# Amino acid usage and tRNA expression underlies proteome adaptation in *Plasmodium falciparum*

**DOI:** 10.1101/2023.03.20.533258

**Authors:** Qian Li, Leonie Vetter, Ylva Veith, Elena Christ, Ákos Végvári, Cagla Sahin, Ulf Ribacke, Mats Wahlgren, Johan Ankarklev, Ola Larsson, Sherwin Chun-Leung Chan

## Abstract

Parasites often depend on metabolic adaptation to the host environment. An adaptive feature of malaria-causing parasites is digestion of hemoglobin (HB) to acquire amino acids (AAs). Here we describe a link between nutrient availability and translation elongation-dependent regulation of gene expression as an adaptive strategy. We show that, unexpectedly, tRNA expression in *P. falciparum* does not match the decoding need as tRNAs decoding AAs that are rare in HB are lowly expressed. This discrepancy renders codons of HB-rare AAs inefficiently decoded and transcripts levels negatively correlated with the requirement of HB-rare AAs for protein synthesis, which are consistent with poor codon optimality causing co-translational mRNA decay. Intriguingly, proliferation-related genes have evolved to require a high level of HB-rare AAs in their encoded proteins, thereby allowing the parasite to control its proliferation by repressing protein synthesis of these genes during nutrient stress. We conclude that the parasite modulates translation elongation by maintaining a discordant tRNA profile as a mechanism to exploit variations in AA-composition among genes as an adaptation strategy.

## Introduction

*Plasmodium falciparum,* a protozoon responsible for malaria, remains a major global health burden. The interactions between this red blood cell (RBC)-infecting parasite and the human host reflect a long-standing arms-race that has driven the evolution of adaptive genetic traits on both sides^1,2^. A notable example is the parasite’s machinery for host HB digestion, which allows the parasite to directly acquire AAs, while obsoleting most AAs biosynthetic pathways^3,4^. Interestingly, the genome of *P. falciparum* is extremely AT-rich (81%) leading to a skewed use of AT-rich codons and their corresponding AAs for protein syntehsis^5^. For instance, AAT and AAA codons account for over 20% of all used codons, and more than 42% of codons decode Asn, Lys, Ile and Leu. This biased AA composition is further reflected by a rapid expansion of low complexity regions in the parasite’s proteome^6,7^. Whether these evolutionary outcomes arise neutrally or are adaptive remains a subject of debate, as conflicting functional studies have failed to establish a consensus^8–10^.

Unlike many unicellular organisms, where tRNA gene copy number correlates with the codon usage^11^, *P*. *falciparum* has a strictly non-redundant set of tRNA genes. Consequently, the most frequently used codons in *P. falciparum* are suboptimal for protein synthesis^12,13^. In some prokaryotes, variation in AA composition can be linked to mRNA and protein expression^14,15^ whereby highly expressed proteins are composed of fewer AAs that are energy-expensive to synthesize^14^. Therefore, although protein function is a strong selection factor on proteins’ AA composition, metabolic constraint may also play a role. This led us to hypothesize that the AA bias in *P. falciparum* may interact with metabolic constraints and thereby influence mRNA and protein expression to drive adaptation; and that this may be particularly important during the intraerythrocytic developmental cycle (IDC) when AAs are largely supplied from digested HB. In this study, we found that AA composition is correlated with mRNA level and protein function during the IDC. Specifically, whereas highly expressed housekeeping genes show a low demand for HB-rare AAs, proliferation-related genes have a high requirement of these AAs for protein synthesis. Furthermore, using a tailored tRNAseq protocol, we observed that tRNAs expression is regulated to exploit translation elongation as a mechanism to modulate genome-wide protein synthesis depending on proteins’ AA composition. These features thereby enable a non-canonical nutrient sensing mechanism for tuning gene expression programs. In summary, the skewed AA usage in *P. falciparum* genome is adaptive to the intraerythrocytic environment.

## Results

### Transcript level is negatively correlated with translation-requirement for amino acids that are rare in hemoglobin

The life cycle of *P. falciparum* involves distinct developmental stages that transit through unique micro-environments within and across hosts. These transitions impose varying metabolic constraints on the parasites. To investigate if the skewed AA usage in the parasite may be adaptive to the metabolic constraints, we asked whether AA composition might influence gene expression across different developmental stages. We therefore quantified transcriptomes using RNA sequencing during all the stages within the RBC and collected published corresponding data from oocyst and sporozoite stages in the vector host^16^. Interestingly, AA usage was sometimes either positively (Gly, Pro, Arg, Val and Ala; Pearson correlation >0.2) or negatively (Ile, Tyr and Asn; Pearson correlation<-0.2) associated with transcript abundance (Fig.1a). Notably, these relationships are strongest during the asexual replicative cycle (ring, trophozoite and schizont). During this cycle, the parasites mature and replicate asexually within RBCs, completed by the release of merozoites that invade new host cells. This replicative cycle is the major determinant of parasite proliferation and biomass increase within the host. A hallmark of these stages is the digestion of host HB to provide AAs^3,4^. Notably, isoleucine, which is the only AA absent in human HB, exhibited the most negative association with transcript abundance in the asexual stage (Fig.1a,b). We therefore hypothesized that transcripts highly expressed during asexual stages have a lower requirement of AAs that are relatively limited in HB (HB-rare), thereby reducing their dependence on external AA availability.

**Figure 1.**
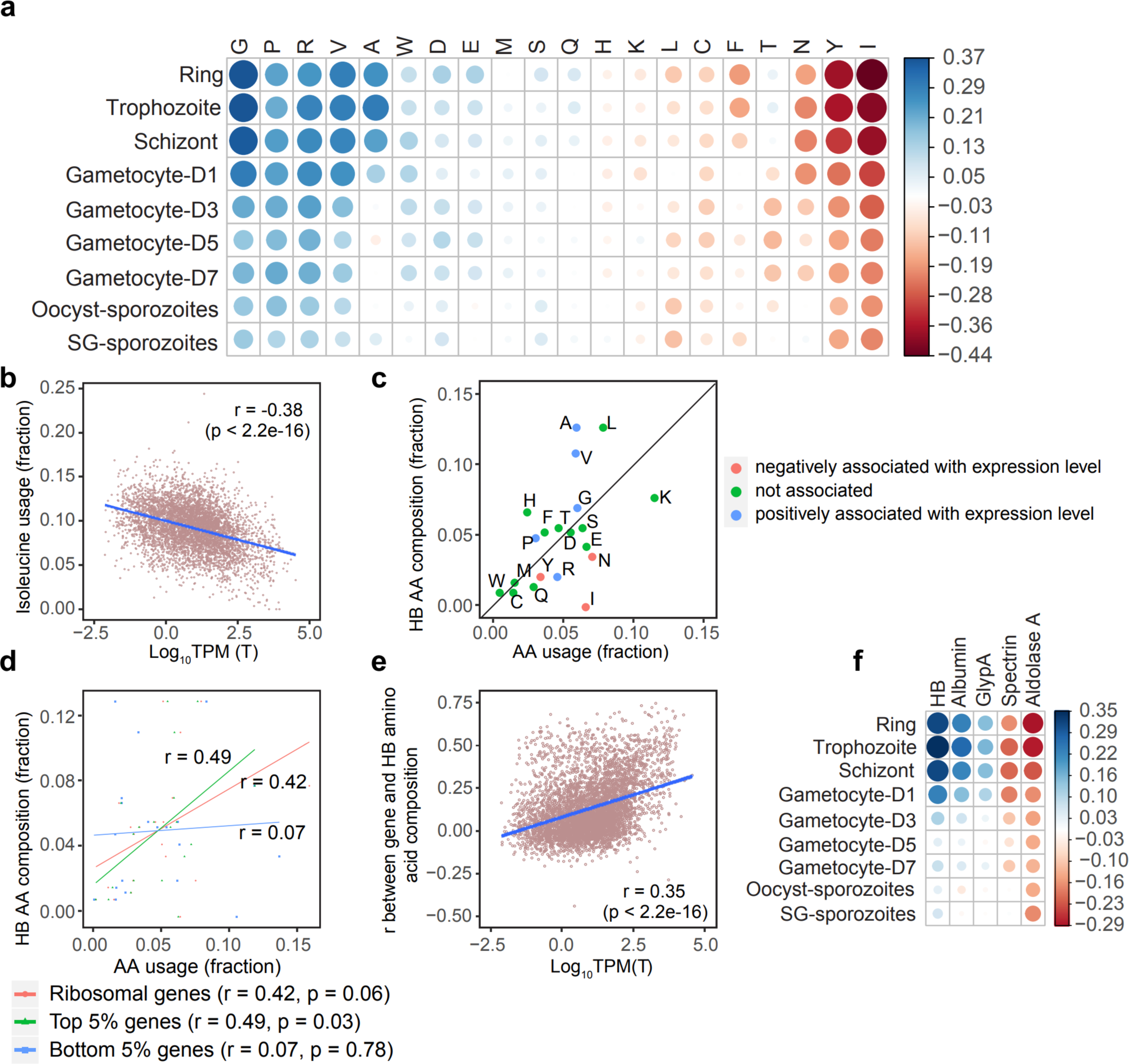
Transcript abundance is associated with the amino acid composition during the IDC. **a,** A representation of Pearson’s correlation coefficient for the frequency of each type of amino acid and transcript abundance (log_10_(TPM)) at different developmental stages. D1-D7 correspond to the number of days post gametocyte induction. **b,** Relationship between genes’ transcript abundance during the trophozoite stage and isoleucine usage. **c,** The relationship between the expression level-weighted requirement of each AA in the trophozoite stage and the composition in host hemoglobin (relative abundance). AAs are colored as negatively or positively associated with mRNA expression if the Pearson correlation (from **a**) was <-0.2 or >0.2, respectively. **d,** Correlation between the AA usage in selected gene subsets and in the relative abundance host hemoglobin. Top and bottom 5% genes were selected following ranking of mRNA expression Log(TPM) values from the trophozoite stage. **e,** The relationship between transcript abundance during the trophozoite stage (x-axis) and the correlation between each gene’s and hemoglobin’s AA usage (y-axis). **f,** Same comparison as (**e**) but showing Pearson correlations coefficients across developmental stages and selected proteins. (HB: hemoglobin; GlypA: Glycophorin A) For all panels, r: Pearson’s correlation coefficient. Coding genes, n=5267. Log_10_TPM values were averaged from n=3.

To test this, we calculated the total AA demand by summing the expression level-weighed AA usage of the transcriptome and compared it to the AA composition of HB (HBA and HBB combined) (Fig.1c). Strikingly, AAs that are negatively correlated with transcript abundance are considered HB-rare when compared to the demand from the transcript pool. Moreover, abundant transcripts show an AA composition more similar to that of HB (Fig.1d-e). This further supports the tenet that abundant transcripts are more dependent on endogenously available AAs during protein synthesis while lowly expressed mRNAs depend on external AA sources. As the positive association is most pronounced during the asexual proliferative stages and is rapidly lost upon sexual differentiation into gametocytes, it may interact with parasite proliferation. Furthermore, this association was stronger for HB AA usage when compared to other RBC-specific proteins or the highly abundant serum albumin protein (Fig.1f), suggesting that the relationship is HB-selective. Interestingly, genes whose AA requirement best matches HB AA frequency are enriched in processes involved in energy homeostasis including glycolysis and also include proteins opposing host immunity, which are both functions essential for survival (Supplementary data 1). On the other hand, genes with the lowest similarity in AA composition relative to HB are often involved in transcription and proliferation (Supplementary data 1). Accordingly, these data suggest that AA bias may enable distinct regulation of survival and proliferation-related genes during nutrient stress.

### Discordance in anticodon to codon pools during asexual stages implicates differential decoding efficiency of amino acids

The association of AA-usage and transcript abundance strongly implicates translation elongation. As *P. falciparum* has a strictly non-redundant set of tRNA genes, a feature that is not shared even by the closely related, yet non-RBC residing parasite, *Toxoplasma gondii* (Extended data Fig.1a), we explored whether the lack of redundancy is compensated through modulating tRNA expression. To this end, we adopted a tRNA sequencing protocol using TGIRT to measure levels of all mature tRNAs (the 45 nuclear tRNAs are referred to as the anticodon pool) together with their aminoacylation levels throughout the IDC (Extended data Fig.1b and Method)^17,18^. We achieved high levels of reads mapping to parasite tRNAs with limited mapping to host tRNAs (from 8.7% in ring to 0.85% in schizont) (Extended data Fig.1c). Moreover, approximately 80% of the sequencing reads mapping to tRNA genes showed complete 5’ to 3’ coverage (Extended data Fig.1d,e). Using unique molecular identifier and assuming that tRNA sequencing is roughly equally sensitive for all tRNAs, the resulting data indicated substantial variation in abundance across anticodons (Fig.2a). For example, the four Ser-tRNAs, which all carry an A-box promoter element divergent from the consensus sequence (Extended data Fig.2a)^19^, are highly abundant (28.7-35.4% of the total anticodon pool). However, despite the cyclic expression pattern observed in most coding transcripts^20^, only minor differences were detected in the anticodon pools across stages (Fig.2a). Similarly, tRNA charging is also stable throughout the IDC (Extended data Fig.2b,c). Finally, despite a distinct transcriptome during each stage (Extended data Fig.2d), the stability of the tRNA pool was matched by a stable tRNA demand as estimated by the codon pool from the transcriptomes (Fig.2b). The stability in both anticodon and codon pools suggests that a transcript’s codon composition may interact with the availability of the required anticodons to regulate gene expression throughout the IDC. Indeed, we observed a discordance between anticodon and codon pools (Fig.2c) where isoacceptors for a specific AA tended to be consistently under- or overrepresented, suggesting that AAs rather than specific tRNAs underlie this property.

**Figure 2.**
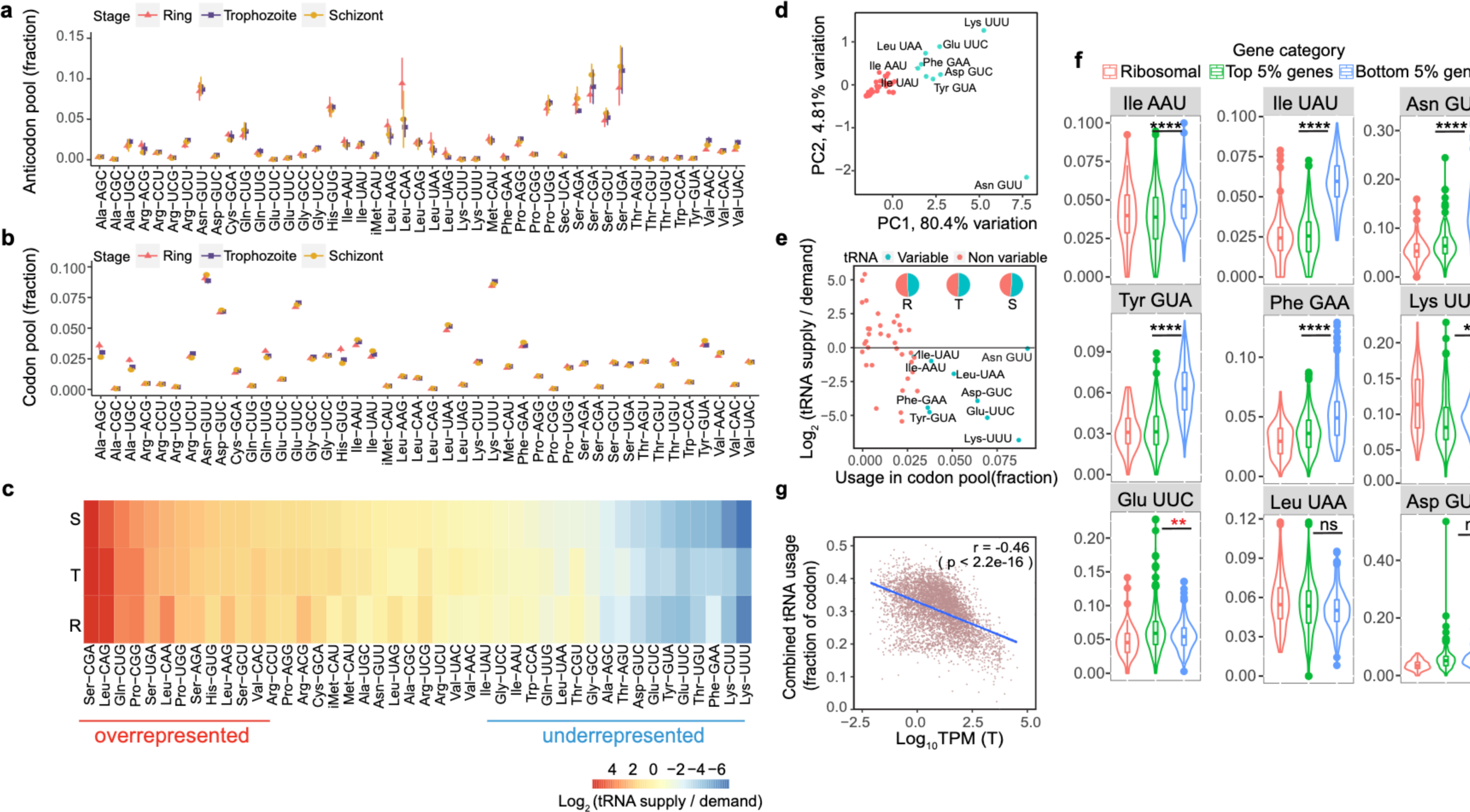
The usage of a subset of tRNAs is associated with transcript abundance. **a,** The relative tRNA supply (anti-codon pool) during the IDC is represented as the number of tRNA sequencing reads for each tRNA relative to the total number of aligned tRNA sequencing reads. **b,** Dynamics of the tRNA demand (codon pool) during the IDC. The tRNA demand of the transcriptome during each stage is represented as the relative expression-weighted (transcripts per million, TPM) codon usage. **c,** Heatmap of averaged log2-transformed ratios of supply to demand (anti-codon/codon) for each tRNA across IDC stages (y-axis: R = ring; T = trophozoite; S = schizont). **d,** Principal component analysis of tRNA demand across all protein coding genes. The tRNAs explaining most of the variance (cyan) were identified and used in subsequent analyses. **e,** A plot of the relationship between the tRNA requirement (codon pool; x-axis) and the log_2_-transformed ratio of supply to demand per tRNA (y-axis) during the trophozoite stage. tRNAs are colored according to their contribution in tRNA variance across genes from (**d**). The line separates regions of underrepresentation (below the line) from overrepresentation (above the line). Pie charts show the combined requirement of these tRNAs from the codon pool. R: Ring; T: Trophozoite; S: Schizont. **f,** A representation of tRNA usage for selected gene subsets (similar subsets as in Fig. 1d). P-values from Mann-Whitney U-test comparing the top and bottom 5% genes are denoted by **** (p<0.0001); ** (p<0.01); ns (not significant, p>0.05). Glu-UUC shows a reversed relationship with higher mean usage in the top 5% genes (red asterisk). **g,** Linear relationship between genes’ mRNA expression during the trophozoite stage and their combined usage demand for Asn-GUU, Ile-AAU, Ile-UAU, Phe-GAA and Tyr-GUA tRNAs. r: Pearson correlation coefficient. For a, b: the mean and SD of n=3 are displayed for each IDC stage.

### Highly abundant transcripts have a reduced decoding-requirement of underrepresented tRNAs

To explore the relationship between the codon-anticodon pools, we used principal component analysis to analyze the pattern of tRNA demand across genes. Nine tRNAs were identified to explain most of the variance in tRNA requirement across genes (Fig.2d). Interestingly, all these tRNAs belong to the previously identified “underrepresented” group that together decode over 50% of the codon pool (Fig.2e). Therefore, despite the high demand, their availability is limited, which could determine the efficiency of translation elongation and thereby affect protein levels. Accordingly, the most abundant transcripts showed a lower demand for six of these tRNAs (Asn-GUU, Ile-AAU, Ile-UAU, Lys-UUU, Phe-GAA and Tyr-GUA) compared to the least abundant transcripts (Fig.2f). This pattern was consistent for transcripts encoding highly abundant ribosomal proteins, with the exception of Lys-UUU. Overall, the combined demand for these tRNAs exhibited a negative correlation with transcript abundance throughout the IDC (Fig.2g). Interestingly, four of these tRNAs exclusively decode asparagine, isoleucine and tyrosine, which aligns with the negative association observed between these AAs and transcript abundance. These findings further support that the tRNAome is coordinated with AA composition to tune genes’ expression.

### Acute amino acid depletion reduces Ile-tRNA availability and targets expression of proliferation-related genes

The negative correlation between translation-requirement for HB-rare AAs and transcript abundance implies AA usage-dependent regulation of protein synthesis upon nutrient stress. To test this, we cultured late stage parasites (32-36 hpi) in AA-depleted medium for 6 hours, which induces acute nutrient stress and forces the parasite to completely rely on AAs from HB. Under this stress, the tRNAome was unaffected but, as expected, the proportion of charged Ile-tRNAs was reduced (Ile-AAU -53% and Ile-UAU -19%) (Fig.3a and Extended data Fig.3a). Interestingly, the reduction in Ile-AAU, which reads Ile codons preferred by highly expressed genes, was more dramatic than that of Ile-UAU (Extended data Fig.3b). This may suggest faster turnover of this tRNA due to translation of highly expressed mRNAs during this window of nutrient depletion. The decrease in charged Ile-tRNAs was not observed when elongation was blocked by halofuginone, an inhibitor of prolyl tRNA synthetase^21^, confirming that the depletion of charged Ile-tRNAs was due to an imbalance between Ile demand and supply during translation elongation (Fig.3a).

**Figure 3.**
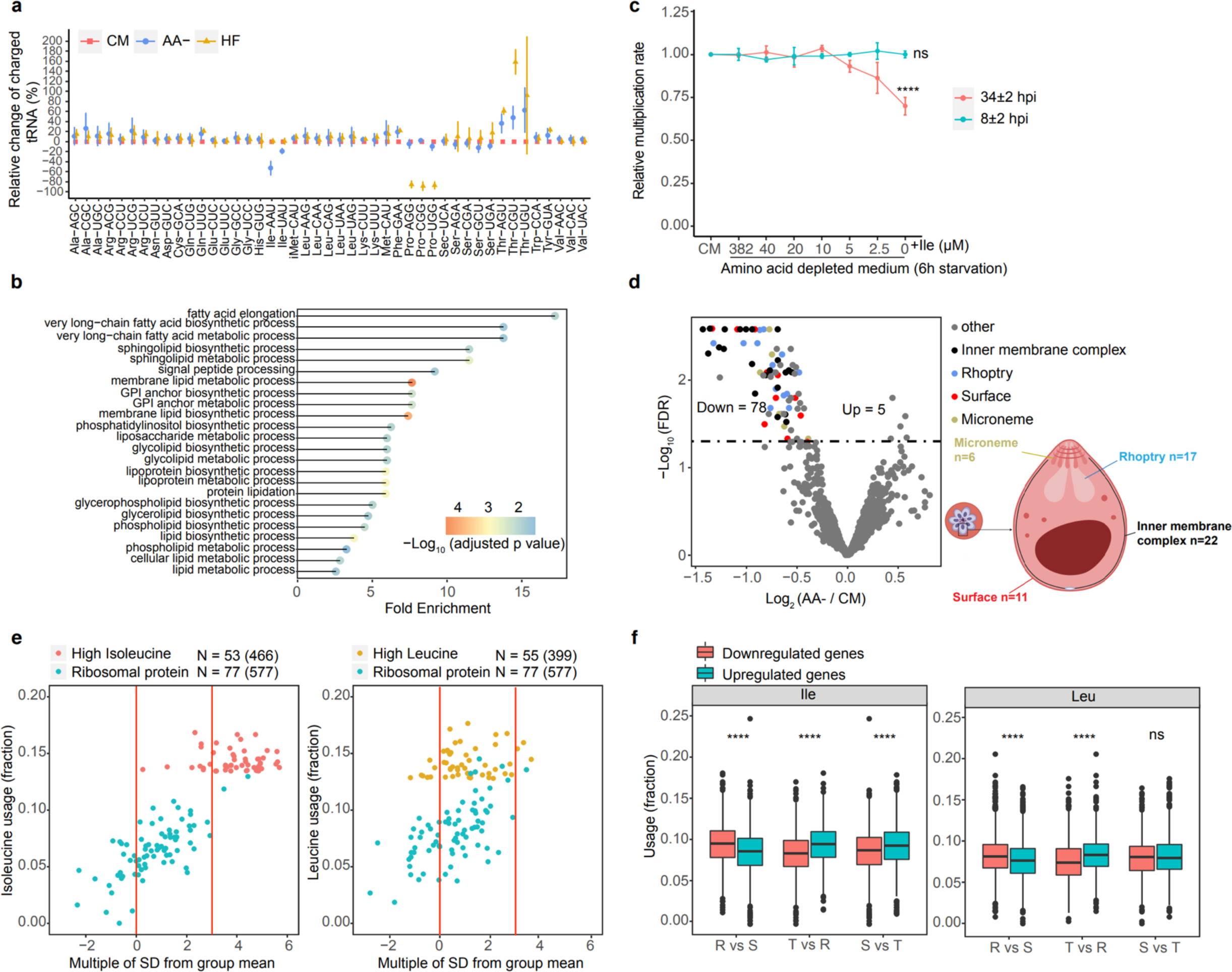
Isoleucine as a sensor of nutrient availability. **a,** Changes of charged tRNA levels in late trophozoite cultures after 6 hours of treatment (CM: complete medium, AA-: AA depletion, HF: halofuginone (70 nM) and AA depletion). Data were normalized to CM for each tRNA separately. Error bars represent SD; n=3 biological replicates. **b,** Gene Ontology (GO) analysis of the top 5% isoleucine-containing genes (n=265), all identified pathways with adjusted p-value <0.05 for the enrichment are shown. **c,** Multiplication rate of parasites after 6 h of AA depletion under varying concentrations of isoleucine. Parasites were starved from either 34±2 hpi (red) or 8±2 hpi (cyan) for 6 h and subsequently recovered in complete medium until all parasites had completed reinvasion. The multiplication rate was normalized to the control group (CM). Error bars represent the SD; n=3 biological replicates. P-values from a two-tailed unpaired t-test are denoted as **** (p<0.0001); ns (not significant, p>0.05). **d,** (left panel) Volcano plot showing 1146 proteins identified in all samples by LC-MS/MS after 6 hours of AA depletion. The dotted line indicates an adjusted p-value=0.05. 56 of the 78 downregulated proteins are associated with the merozoite stage, and their reported localizations are indicated (right panel). **e,** A plot showing the deviation (number of SDs) of isoleucine (left) and leucine (right) usage in selected *P. falciparum* proteins relative to usage in orthologous proteins. High isoleucine or isoleucine subsets contain the 5% of proteins with the highest frequency of the respective amino acid while the ribosomal proteins subset is the same subset as in Fig. 1d. Only genes with >100 orthologues were considered as functionally conserved and analyzed. Numbers in brackets indicates average orthologue counts. Red lines indicate 0 and 3 SD. **f,** Box plot showing isoleucine (left) and leucine (right) usage in up- and downregulated genes during stage progression (R = ring, T = trophozoite, S = schizont stage). P-values from Mann-Whitney U-test are indicated with **** (p<0.0001); ns (not significant, p>0.05).

As acute AA depletion only reduces the availability of charged Ile-tRNAs, genes with high Ile usage would be expected to be selectively affected. Gene ontology (GO) analysis showed an enrichment of various lipid metabolic pathways among the 5% genes with the highest Ile content (Ile-rich) (Fig.3b). In contrast, genes with the highest leucine content, which is also decoded by AT-rich codons, show no pathway enrichment at FDR<0.05. As lipid metabolic pathways are essential for membrane biogenesis during schizogony and merozoite production^22,23^, this may affect parasite proliferation. Indeed, an experiment where ring or trophozoite parasites were cultured in AA-depleted medium for 6 hours followed by recovery in complete medium for 48 hours showed that only treated trophozoite cultures had reduced multiplication rate in the following cycle (Fig.3c). This could be rescued, in a dose-dependent manner, by supplementing with Ile during AA-depletion (Fig.3c). Treated ring cultures were not affected because proliferation has not commenced at this stage. Additionally, quantitative proteomic analysis revealed that 56 out of the 78 proteins (71.8%) downregulated upon trophozoite starvation were merozoite-related proteins, these are predominantly associated with membrane and organellar compartments, particularly the inner membrane complex (Fig.3d and Supplementary table 2). This indicates that proliferation-related Ile-rich genes are targeted upon nutrient stress.

To address whether gene-specific high Ile usage has evolved as part of an adaptation strategy, we analyzed Ile usage in orthologues of Ile-rich genes. Strikingly, over 80% (43/53) of the conserved Ile-rich genes (defined as having >100 orthologues) encode an Ile content that is >3 standard deviations higher than the mean Ile content in the respective orthologue group (Fig.3e)^24^, suggesting that the high Ile content in Ile-rich genes is unlikely to reflect conservation of protein function, which should universally preserve high Ile content in the orthologues. Indeed, this was the case when assessing leucine-rich proteins (only 3/55 genes have a leucine content >3 S.D. than the mean). This analysis therefore supports that the high Ile content in Ile-rich genes is adaptive and serves mainly to tune their expression.

To further assess the relationship between Ile usage and gene expression, we considered the delayed stage progression phenotype, described upon prolonged AA-starvation^25,26^. We confirmed the stage-delay phenotype even with short-term AA-depletion (Extended data Fig.4a). During ring-to-trophozoite and trophozoite-to-schizont transitions, upregulated transcripts exhibited a higher Ile-content than downregulated transcripts. Conversely, the opposite pattern was observed during schizont-to-ring transition (Fig.3f and Extended data Fig.4b,c). Additionally, transcripts that displayed the highest variability throughout the IDC encoded a higher Ile content compared to those with the lowest variation between stages (Extended data Fig.4d). This analysis therefore provides a rationale for the stage-delay phenotype and explain the higher sensitivity to Ile withdrawal reported in pre-S phase compared to post-S phase parasites^26^. Importantly, it also implicates a role of the across-gene variation in AA composition in modulating stage progression.

### High Ile usage leads to ribosome stalling in absence of mRNA degradation during nutrient stress

While our results indicate that variation in AA composition affects protein synthesis via tRNA-dependent modulation of translation elongation, the link to mRNA level remained unclear. A plausible mechanism is induction of co-translational mRNA decay following ribosome stalling^27–29^. As acute AA-depletion reduces Ile-tRNA charging (Fig.3a), this model allows us to delineate the relationship to ribosome stalling and mRNA levels. As indicated by polysome fractionation, there was a reduction in polysome formation with a concomitant increase in 80S monosomes after AA-depletion (Fig.4a), which is expected following suppression of translation via increased eIF2α phosphorylation during AA starvation^30,31^. To determine how individual mRNAs were regulated upon AA starvation, we quantified polysome-associated mRNA using RNA sequencing. We first compared Ile usage between the 247 and 152 transcripts that showed increased and decreased polysome-association, respectively, upon AA-depletion (Extended data Fig.5a). This revealed that transcripts with increased polysome association decode Ile more often than those with decreased polysome association (Fig.4b). This contrasted with our expectation that lower Ile-tRNA level would promote co-translational decay of transcripts with higher Ile-requirement for their translation. Instead, we observed a strong correlation between the change in total and polysome-associated mRNA levels among these transcripts (Fig.4c). However, we also found that whereas changes in polysome-associated mRNA and protein levels showed a substantial correlation for transcripts with decreased polysome association (the larger subset with a log_2_ fold-change <0 for polysome-associated mRNA), this correlation was lower for transcripts with increased polysome association (Fig.4d). This suggests that the increase in ribosome loading reflects ribosome stalling without reduced mRNA level. This is also consistent with an increased association of mRNA with polysomes without augmented protein levels.

**Figure 4.**
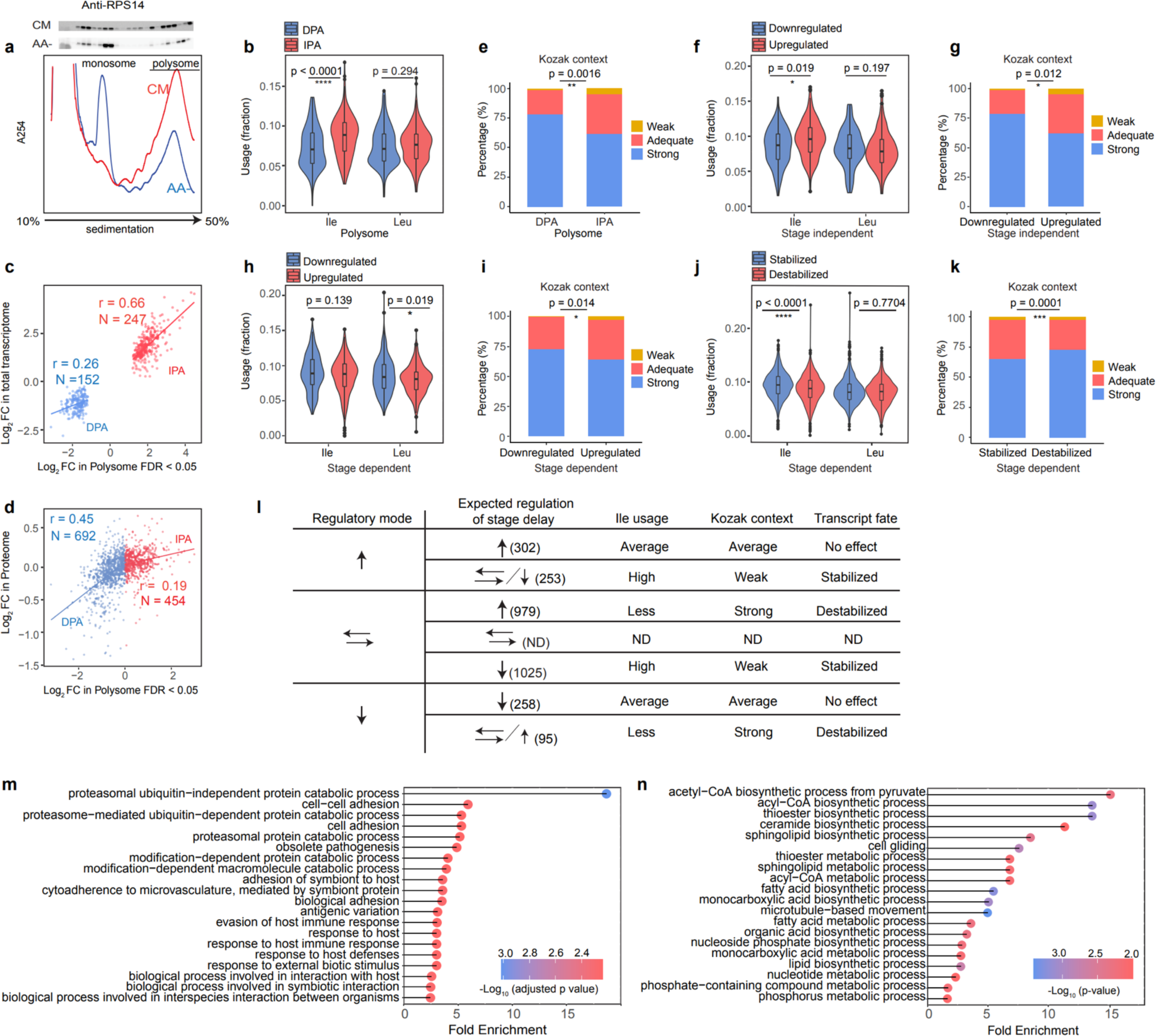
Polysome profiling reveals targeted reprogramming after AA starvation. **a,** (lower panel) Representative polysome tracings across a 10-50% sucrose gradient from parasites growing in complete medium (CM) and AA-depleted medium (AA-). Fractions containing monosomes and polysomes are indicated. (upper panel) Immunoblotting of each fraction against RPS14. **b,** Differences in isoleucine and leucine usages between genes with decreased (DPA) or increased (IPA) polysome association. **c,** Correlation of changes in polysome-association and total mRNA separately for up- or down-regulated genes. Transcripts with an FDR<0.05 and log_2_ fold-change <-1 or > 1 from the analysis of polysome association are shown. **d,** Correlation of changes in transcripts’ polysome association and changes in corresponding protein levels separately for up- or down-regulated genes. **e,** Kozak context of genes with decreased (DPA) or increased polysome association (IPA). **f,g,** Stage independent differences in (**f**) isoleucine and leucine usage; (**g**) Kozak context for stage-independent up- or downregulated genes. **h,i,** Stage dependent differences in (**h**) isoleucine and leucine usages; (**i**) Kozak context for stage-dependent up- and downregulated genes. **j,k,** Stage dependent differences in (**j**) isoleucine and leucine usage; (**k**) the Kozak context in transcripts that remained stable after amino acid depletion but were expected to be downregulated (stabilized) or upregulated (destabilized) according to the categorization shown in Extended data Fig.5d. **l,** A summary table of the transcriptome changes and associated transcript features after 6 hours of AA depletion. ↑ : increased transcript abundance, ↓ : decreased abundance, ⇄ : unchanged abundance. ND: not determined. **m,n,** Gene Ontology analysis of stage-independent (**m**) downregulated and (**n**) upregulated genes. Only 20 significant terms with the highest enrichment are shown. Kozak context - strong: RnnATGR; adequate: RnnATGY/YnnATGR; and weak: YnnATGY (ATG= start codon). Statistical significance was assessed using Mann-Whitney U test in (b), (f), (h), (j); and Fisher’s exact test for the Kozak context analysis in (e), (g), (i) and (k). ****(p<0.0001); **(p<0.01); *(p<0.05)

**Figure 5.**
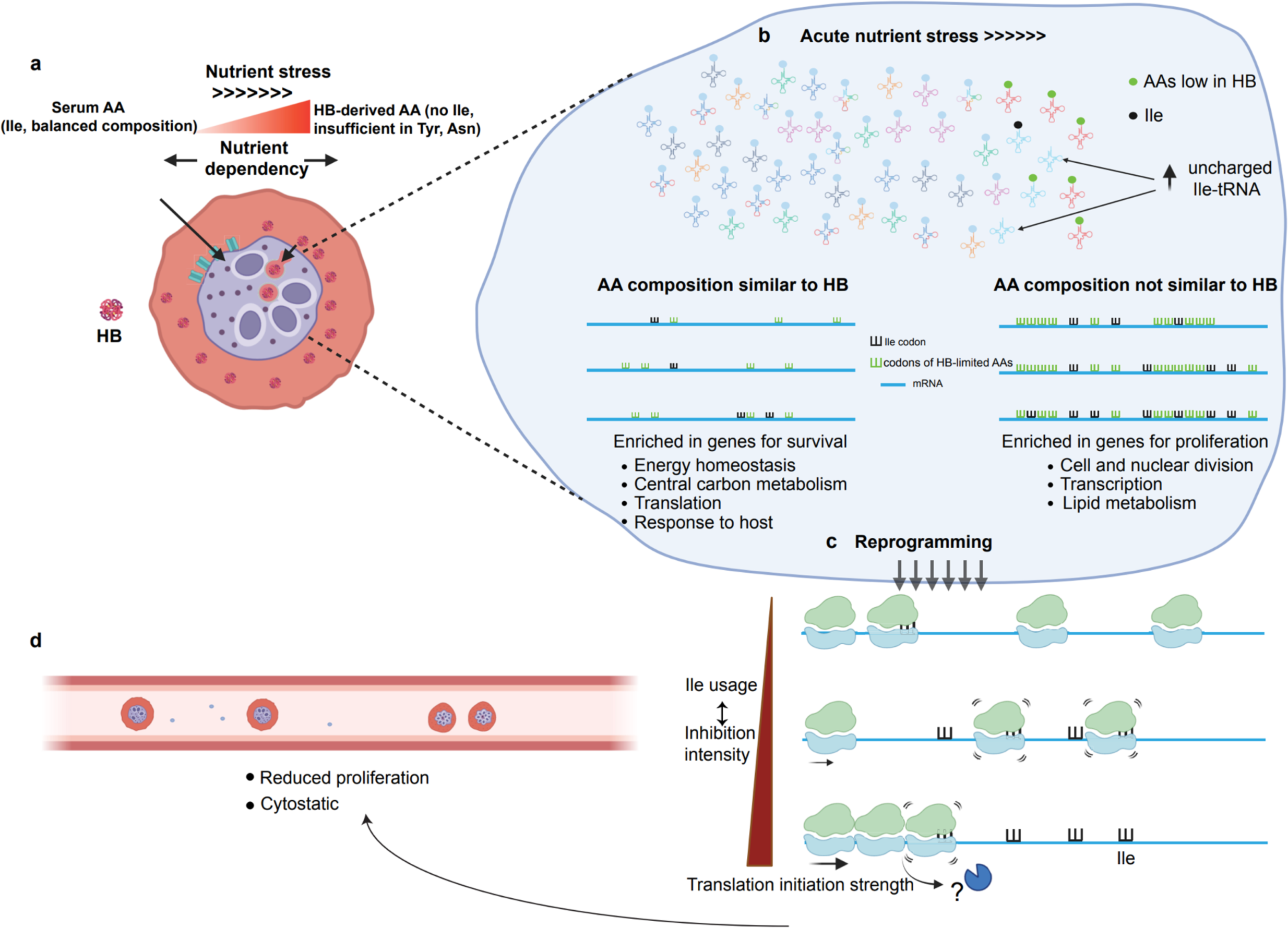
Proposed model of a nutrient-sensing mechanism in *P. falciparum*. **a,** *P. falciparum* has two major amino acid acquisition pathways to aid its parasitic lifestyle within RBCs. Most amino acids can be predominantly supplied through digestion of host hemoglobins (HB), but a few amino acids, in particular isoleucine, need to be imported exogenously from host serum to meet the cellular requirement. **b,** Nutrient deprivation reduces the accessibility to exogenous nutrient sources, prompting the parasite to rely more heavily on HB-derived AAs. A reduction in serum Ile level causes an accumulation of uncharged Ile-tRNAs and a selective decrease in translation. Specifically, proteins with housekeeping functions have evolved to depend primarily on the stable supply of AA from host HB, thereby minimizing their reliance on external nutrients. This adaptation allows for translation of these proteins also under acute nutrient fluctuation. In contrast, proliferation-related genes are targeted due to their high Ile usage. **c,** The changes in ribosome loading also reprogram the transcriptome. Differential stabilization of transcripts depends on the interplay between translation initiation and Ile-codon-induced stalling. **d,** Targeted translation repression of proliferation-related genes and transcriptome reprogramming autoregulate parasite proliferation and drive the cell into a cytostatic “hibernation” state.

Ribosome collisions driven by excessive ribosome stalling can trigger the conserved No-Go-Decay (NGD) pathway, leading to degradation of the transcript^32,33^. Accordingly, the balance between ribosome stalling and collision will depend on translation initiation and the time needed to resolve the stalling event^34^. We therefore analyzed if the Kozak context, which is a major determinant of initiation strength, associated with the change in polysome association upon AA-depletion. This analysis suggested that mRNA with increased polysome association had a weaker Kozak contexts than mRNA with decreased polysome association (Fig.4e and Extended data Fig.5b)^35^. Therefore, some high Ile-decoding mRNAs may have prevented ribosome collision by having a less efficient initiation and thereby escape NGD during AA-depletion. Furthermore, these observations suggest that this interplay may instead lead to Ile-dependent stabilization of transcripts and thereby contribute to a transcriptome priming during AA-depletion, a mechanism that would reduce the requirement for re-synthesis of targeted mRNAs upon stress relief.

### The interplay between translation initiation and ribosome stalling determines transcript fate

We determined changes in total mRNA levels using samples collected in parallel with the polysome-associated RNA samples and identified 555 and 353 genes with increased and decreased mRNA levels, respectively, after AA-depletion (Extended data Fig.5c). As AA-depletion delayed stage progression, we assigned differentially expressed genes as either stage dependent or stage independent by comparing to gene expression differences between trophozoite and schizont stages (Extended data Fig.5d). Among the stage-independent changes, higher Ile usage and a weaker initiation context were observed in transcripts with increased mRNA levels compared to those with decreased mRNA levels (Fig.4f,g and Extended data Fig.5e). In contrast, there were no differences in the Ile usage and only a minor difference in initiation context when comparing stage-dependent subsets (Fig.4h,i and Extended data Fig.5f). As these features were shared when assessing polysome-associated mRNA and stage-independent total mRNA changes it supports the tenet that that mRNAs were stabilized following increased polysome association. We further assessed this by studying mRNAs that were not regulated as expected due to delayed stage progression (Extended data Fig.5d). The motivation being that these mRNAs may have been either “stabilized” (i.e., they were expected to be downregulated but were unchanged upon AA starvation) or “destabilized” (i.e., they were expected to be upregulated but were unchanged upon AA starvation) during reprogramming. Indeed, the stabilized transcripts showed slightly higher Ile usage and weaker initiation context than the destabilized transcripts (Fig.4j,k and Extended data Fig.5g). As these features are associated with increased polysome association it is consistent with a stabilizing effect for transcripts with increased polysome-association and reduced elongation. Interestingly, many transcripts encoding ribosomal and proteasomal proteins were found in the subset of stage-independent downregulated genes (Fig.4m and Supplementary data table3). These changes may lead to reduced proteome turnover and allow the parasites to cope with prolonged AA-starvation^25^. In contrast, upregulated stage-independent genes were enriched in various metabolic pathways (Fig.4n and Supplementary data table4). As upregulation may depend on transcript stability but not increased protein output, these changes may prime a cell for rapid metabolic reactivation after nutrient stimulation.

Additionally, we used anota2seq to further explore the relationship between polysome-associated and total mRNA (Extended data Fig.6a). Anota2seq identifies changes in polysome-associated mRNA occurring independent of changes in total mRNA (denoted “translation”); congruent changes in total and polysome-associated mRNA (“abundance”); and changes in total mRNA not reflected by corresponding changes in polysome-association (“buffering”)^36^. Anota2seq analysis revealed many buffered mRNAs upon AA starvation. Notably, buffered mRNAs with increased (“buffered, mRNA up”) or decreased (“buffered, mRNA down”) total mRNA levels also correspond to mRNAs with relatively decreased or increased ribosome association, respectively. Buffered transcripts therefore exhibit a discordance between ribosome association and total mRNA levels and would be expected to show differed RNA features compared to those with altered polysome association. Indeed, the “buffered, mRNA down” subset shows higher Ile usage than the “buffered, mRNA up” subset, yet having stronger Kozak initiation context (Extended data Fig.6b-d). Tentatively, we reason that despite the stalling of ribosomes on Ile-codons, the “buffered, mRNA down” subset was destabilized due to strong translation initiation, which lead to high ribosome influx and collisions. Accordingly, we propose a model for how a global reprogramming can be achieved during nutrient stress (Fig.4l & 5).

As stalling at Ile codon is instrumental to the observed transcriptome reprogramming, we believe that the natural expansion of homo-Ile repeats, which present strong stalling motifs, would be under purifying selection. Without functional constraint, AA repeats often appear at a higher frequency than expected by chance due to replication slippage^37^. This is true for most AAs, as observed by the higher frequency of most homo-di-amino acid motifs than expected in *P. falciparum* genome. In contrast, di-Ile occurs less frequently than expected (Extended data Fig.7). Only di-methionine motif exhibits the same trend, yet methionine is under functional restraint as the initiatior AA. Furthermore, longer stretches of Ile also occur at a lower frequency than expected, which are opposite to other hydrophobic AAs. Therefore, occurrence of Ile repeats is likely under non-neutral selection, further asserting a central role for Ile in stress-mediated reprogramming.

### Adaptation-associated bias in amino acid usage may be conserved

From an evolutionary perspective, this study provides evidence that metabolic constraints can be a significant driving force in the evolution of a protein’s primary sequence. As a proof-of-concept we compared the AA compositions of proteins in glycolysis and the pentose phosphate pathway (PPP) to the AA composition of HB in several organisms with different genome GC-contents. While both pathways are highly conserved and utilize glucose-6-phosphate as input, glycolysis generates ATP, while PPP supplies ribose-5-phosphates for nucleic acid synthesis^38^. Therefore, despite competing for the same nutrient flux, glycolysis is essential for energy homeostasis, while PPP favors proliferation. Importantly, our results indicate that glycolysis proteins in all investigated *Plasmodium* and *Babesia* spp., parasites that replicate within RBCs^39^, exhibit a stronger correlation to host HB compared to components of the PPP (Fig.6). This supports the notion that these organisms might have exploited the across-gene variation in AA composition to adapt to metabolic constraints. Interestingly, despite also residing in RBCs, *Theileria* spp. does not exhibit this pattern (Fig.6). As schizongy in *Theileria* exclusively occurs in monocytes^40^, not RBCs, this replicative mode may have reduced HB-dependence and its associated adaptive selection pressure. Given the close lineage relationship between *Babesia* and *Theileria,* the contrasting observations between the two genera indicate that this type of selection is highly dynamic and depends on the host metabolic constraints.

**Figure 6.**
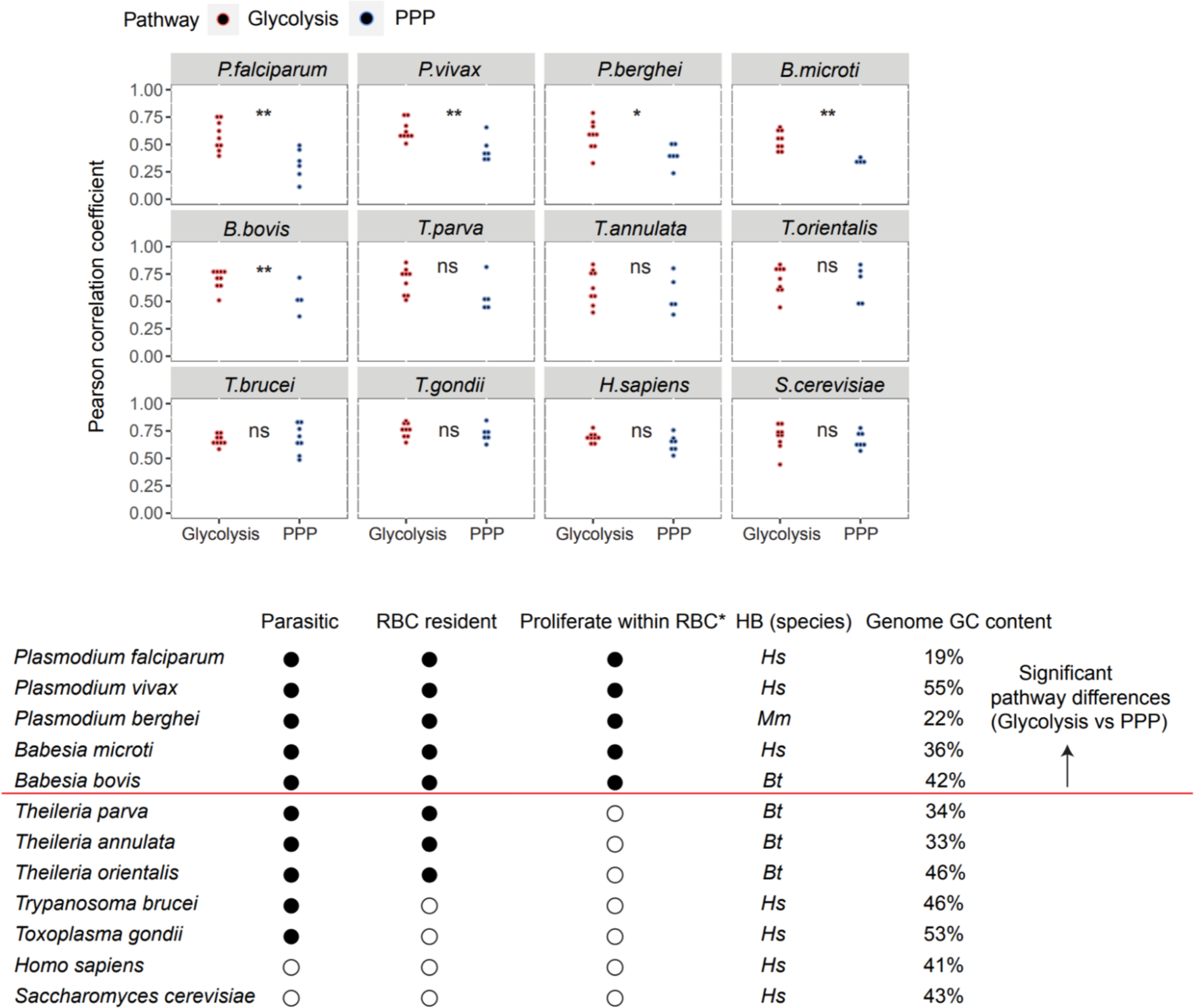
Lower dependence on external nutrient sources is conserved in housekeeping genes. (Upper panel) Correlations between AA composition of host hemoglobin (HB) and enzymes involved in glycolysis or the pentose phosphate pathway (PPP) in various species. (Lower panel) Overview of assessed species. Filled circle: Yes, unfilled circle: No. Proliferate within RBC* indicates whether proliferation within RBC constitutes the major determinant of biomass increase within the host. HB (species) specifies the host species used for HB correlations. *Hs*: *Homo sapiens*, *Mm*: *Mus musculus*, *Bt*: *Bos Taurus*. The red line separates species (above the line) showing significant pathway differences in AA composition. P-values from Mann-Whitney U-test are indicated by ** (p<0.01); * (p<0.05); ns (not significant, p>0.05).

## Discussion

This study reveals translation elongation as a prominent mechanism to tune gene expression at a transcriptome-wide level. Despite considerable recent progress, gene regulation mechanisms utilized by the parasite remain largely elusive. Transcriptional control is believed to play a lesser role as compared to other eukaryotes due to the paucity of specific transcription factors, whereas a significant enrichment of RNA-binding proteins points towards an elaborated post-transcription regulatory network^41^. Using TGIRT-seq to study the tRNAome, we demonstrate that tRNAs are expressed in discordance with the decoding requirement of the transcriptome. This seems to have a role in enabling a systemic modulation of protein synthesis during the translation elongation step. Dynamic alteration of tRNA availability as a regulatory mechanism has been implicated in cell-fate decision, tissue-specification and pathological conditions, as well as during changes in growth condition and stress levels^42–51^. Various mathematical models and *in vitro* studies have also recapitulated the effect of tRNA availability in determining ribosome flux and protein synthesis rate^31,34,52^. Notably, the average length of *P. falciparum* coding-sequences is substantially longer than in other eukaryotes^5^. This emphasizes the relative importance of tRNA-mediated regulation. Discordant tRNA expression may also raise the prospect of adaptive translation, which exploits translation infidelity to diversify the proteome^53–55^. Notably, experimentally manipulated tRNA mis-expression is associated with adaptive genome instabilities in yeast^56–58^, emphasizing the prominent influence of tRNAs during the evolution of genome.

Our study unraveled that the disparity in decoding dynamics and transcript abundance significantly associated with the AA identity. Similar findings were described in other organisms^27,59,60^, but mostly considered as an associated effect that is secondary to the protein primary sequence. Yet in *P. falciparum*, AAs that are rare in HB are most negatively associated with gene expression level. AAs sufficiently supplied from HB digestion are expected to be more stably available than AAs that need to be acquired exogenously from the serum, as their availability may be affected by e.g. the host’s feed-fast cycle, malnutrition and the progression of the infection itself ^61–63^. Thereby, evolving an AA composition similar to HB ensures stress-insensitive expression of proteins with essential housekeeping functions. This argues that the across-gene variation in AA composition is strongly shaped by metabolic constraints and reflects metabolic adaptation. Therefore, this study provides the unique perspective that AA identity can be selected primarily on the basis of decoding dynamics, rather than protein function.

*Plasmodium spp.* lacks the conserved mTOR pathway, which is a critical node modulating translation^64^. Moreover, the parasite can mount an effective response and recovery from AA starvation in absence of phosphorylatable eIF2α^25^, which is a key component of the integrated stress response (ISR). Therefore, additional stress response mechanisms have been expected to exist. Here we show that variation in AA composition may represent one such mechanism by efficiently partitioning genes’ responsiveness upon nutrient stress. Similar to ISR, this allows a transcriptome-wide response to be mounted in a rapid, targeted and graded fashion. This mechanism is highly robust and adept as, unlike canonical stress responses, it does not rely on kinase-mediated signaling cascades, thereby allowing decentralized allocation of resource based solely on demand and supply.

The high AT content of the *P. falciparum* genome is maintained by a significant excess of G:C to A:T transition *in vitro*, prompting the speculation that such mutational bias contributes to adaptive evolution^65^. Coincidentally, AAs that are negatively associated with gene expression are invariably decoded by AT-rich codons. It is compelling to suggest that the substitution to and insertion of these AAs may be continuously selected during growth, as these may help to optimize resource reallocation and increase fitness. To some extent, this trait of metabolic adaptation can partially account for the compositional bias in the proteome and the AT-biased mutational rate, at least within the coding region.

Contrary to the conventional belief that functional selection and neutral drift are the main drivers of protein evolution^66,67^, our study implies that even a global selection sweep that affected the primary sequences of many proteins may neither be functionally relevant nor neutral in its nature. This adds new perspectives to the understanding of genome and protein evolution including adaptive mutation bias and frequent occurrence of low complexity regions in *P. falciparum* genome^6,37,66^.

## Methods

### Parasite culturing

*Plasmodium falciparum* strain NF54 were cultured with 4% human O+ erythrocytes hematocrit in RPMI 1640 medium (Thermofisher Scientific) supplemented with 10% A^+^ human serum (Karolinska Hospital blood bank, Stockholm, Sweden), 50 μg/L gentamicin (Gibco) and 2mM L-glutamine (Hyclone). Cultures were maintained at 37°C in 90% N_2_, 5% O_2_, 5% CO_2_ gas mixture on a 50-rpm shaker. Parasites were maintained at <3% parasitemia and routinely synchronized by treating with 5% sorbitol solution (Sigma) for ring stage and by using percoll gradient (Cytiva) for late stage.

### Gametocytes induction

The gametocyte-producing NF54 Peg4-tdTomato transgenic line was used for the production of gametocytes in this study^68^. Prior to induction, parasites were cultured at normal growth condition described above, except that the cultures were individually gassed with 96% N_2_, 1% O_2_, and 3% CO_2_ and maintained at ≤ 1% parasitemia. The parasitemia was monitored every 48 hours before the initiation of gametocyte induction. Induction was performed at 10-12% parasitemia using the commitment assay described in Fivelman *et al*^69^. From day 0 onward, gametocytes were maintained with N-acetyl glucosamine in order to inhibit asexually replicating parasites in the cultures.

### RNA isolation

For samples collected during the IDC, cultures were expanded to 30 mL at 3% hematocrit and ∼8% parasitemia, 10 mL culture were collected for RNA extraction at each time point (8-12, 24-28 and 40-44 hpi). For samples collected during gametocytogenesis, 10mL cultures were collected for RNA extraction at 1, 3, 5 and 7 days after gametocyte induction.

#### For tRNA library

Small RNA (<200 nt) was isolated directly from the pellets of infected RBCs using the *mir*Vana miRNA isolation kit (Invitrogen) according to the manufacturer’s instructions. 600 μL lysis buffer was added to 150 μL infected RBCs and the cell suspension was vortexed vigorously to completely lyse the cells. 1/10 volume of miRNA Homogenate Additive was added to the cell lysates and the samples were incubated 10min on ice. After addition of one v/v ratio of Acid-Phenol: Chloroform to the lysate, the samples were vortexed vigorously and centrifuged at 10,000 g for 5 min and the aqueous phase was transferred to a new tube. 1/3 volume of 100% ethanol was added to the aqueous phase and subsequently passed through the column. The flow through was mixed with 2/3 volume of 100% ethanol and passed through a second filter. Small RNAs were immobilized on the filter. Two consecutive washing were performed using miRNA wash solution 1 and wash solution 2/3, respectively. Small RNAs were eluted in 70μL pre- heated (95°C) nuclease-free water.

#### For mRNA library

Total RNA was extracted from 100 μL infected RBCs using Trizol reagent (Sigma). 1 mL Trizol was added to the cell pellet and pipetted up and down until complete homogenization of the lysate. The protocol was then carried out according to the manufacturer’s instructions. Total RNAs and small RNAs were quantified by Qubit RNA high sensitivity kit (Invitrogen).

### tRNA library construction

#### Periodate oxidation and β-elimination

RNA oxidation and β-elimination were performed as previously described^18^. 2 μg isolated small RNA was treated in 100 mM NaOAc/HOAc, pH 4.8 and freshly prepared 50 mM NaIO_4_ at room temperature for 30 min. The reaction was quenched by incubating with 100 mM glucose for 5 min at room temperature. The treated RNA was purified first with RNA clean-up and concentration micro-elute kit (Norgen Biotek Corp.), and then by ethanol precipitation. For β-elimination, the purified RNA was treated in 60 mM sodium borate, pH 9.5 at 45°C for 90 min. Subsequently, the RNA was purified with RNA clean-up and concentration micro-elute kit followed by ethanol precipitation. Two short RNA standards with A-end and C-end were mixed in equimolar ratio and added to the purified RNA to a final concentration of 0.4 pmol/μg small RNA).

Two synthetic RNA standards:

A-ending: 5’-GCAGAUGGCUUCAAUUGCUAUUAAGGACCA

C-ending: 5’-GCAGAUGGCUUCAAUUGCUAUUAAGGACAC

#### tRNA library generation

This was carried out as previously described with minor modifications^17^. For cDNA synthesis, optimized R2R DNA (1:1 ratio of T-ending and G-ending oligos) was annealed to complementary R2 RNA in 100 mM Tris-HCl, 0.5 mM EDTA, pH 7.5 at 82°C for 2 min, then cooled down to 25°C with a ramp rate of 0.1°C /s^70^. A total of 100 ng tRNA (≈4 pmol) was mixed with 4 pmol primer mixture and pre-incubated at room temperature for 30 min with low salt buffer (50 mM Tris-HCl, pH7.5, 75 mM KCl, 3 mM MgCl_2_), 10 mM DDT and 500 nM TGIRT-III enzyme (InGex, Inc). dNTPs mix were added to a final concentration of 1 mM to initiate the reaction. The template switching reaction was performed at 42°C for 16 h.

The reactions were terminated by adding 1 μL of 5 M NaOH and incubated at 95°C for 3 min. The reactions were neutralized by addition of 1μL of 5M HCl. cDNAs were then purified by MinElute Reaction Cleanup Kit (QIAGEN) and eluted in 10 μL nuclease free water. 10 μM R1R DNA containing UMI sequences was adenylated with 2 μL of Mth RNA ligase (New England Biolab) at 65°C for 1 h and subsequently inactivated at 85°C for 5 min. 2 μM of the adenylated R1R DNA was ligated to the purified cDNAs at 65°C for 2 h with Thermostable 5’ AppDNA/RNA ligase (New England Biolab). After inactivation at 90°C for 3 min, the ligated cDNAs were purified with MinElute Reaction Cleanup Kit (QIAGEN) and eluted in 23 μL nuclease free water. PCR amplification for Illumina sequencing was performed using 2X Phusion High-Fidelity PCR Master Mix (Thermofisher Scientific) for 12 PCR cycles (98°C 5s, 60°C 10s, 72°C 10s) using R1 multiplex PCR primer and barcoded primer. 1.3X Agencourt AMPure XP beads (Beckman Coulter) were used to clean up the libraries and get rid of primer dimers before Illumina sequencing. The quality and quantity of each library was analyzed on the Bioanalyzer with a High Sensitivity DNA analysis Kit (Agilent technologies) and Collibri Library Quantification Kit (Invitrogen).

*R2 RNA:*

5’-GAUCGGAAGAGCACACGUCUGAACUCCAGUCAC-3SpC3/

*R2R DNA T-ending/ G-ending:*

5’-GTGACTGGAGTTCAGACGTGTGCTCTTCCGATCT/G

*R1R DNA:*

5’-/5Phos/GATANNNNNNNGATCGGAAGAGCGTCGTGTAGGGAAAGAGTGT/3SpC3/

*R1 multiplex PCR primer:*

5’-AATGATACGGCGACCACCGAGATCTACACTCTTTCCCTACACGACGCTCTTCC GATCT

*Barcoded primer:*

5’CAAGCAGAAGACGGCATACGAGAT[BARCODE]GTGACTGGAGTTCAGACGTGT GCTCTTCCGATCT

### mRNA library construction

100 ng total RNA was used to construct mRNA library using TruSeq Stranded mRNA kit (Illumina). The protocol was carried out following manufacturer’s instruction. The quality and quantity of each library was analyzed on the Bioanalyzer with a High Sensitivity DNA analysis Kit (Agilent technologies) and Collibri Library Quantification Kit (Invitrogen).

### Library sequencing and analysis

#### tRNA library read processing

All libraries were sequenced on an Illumina Nextseq 550 platform with a single-end read length of 120 bp. Base calling and demultiplexing were performed with Generate FASTQ (v2.0.1). Unique molecular identifiers (UMI) were extracted using UMI_tools (v1.1.2)^71^. Adapter sequences were clipped from the reads using fastx toolkit (v0.0.14) with a size filtering to discard reads <15bp. These analyses were performed on the galaxy platform.

For our system, we expect that the low complexity tRNA transcriptome of the parasite would generate the majority of reads sequenced in the libraries, with the few remaining reads originated from the highly complex human tRNA transcriptome as well as from other non-tRNA small RNA species. Therefore, a combined tRNA reference (including Plasmodium nuclear and apicoplast tRNAs and all human tRNAs from GtRNAdb^72^) was first compiled and curated using sequence data generated from a pilot sequencing trial. The curation includes re-defining the exact 5’ and 3’ position of all tRNAs, addition of 3’-CCA to all tRNAs, changing the reference from A to G for tRNAs that has inosine modification on position 34 and the addition of a 5’ G base on His-tRNA^73^.

The sequence reads were then aligned to the curated tRNA reference using bowtie2 (v2.2.1) with sensitive local mode^74^. The unaligned reads were then aligned to the 3D7 genome (v. PlasmoDB-46) with sensitive local mode to identify mapping to other small RNA species. Two addition steps were performed to assess the outcome of the mapping. First, the unaligned reads from the genome mapping were again aligned to the tRNA reference with very sensitive local mode, which should yield very few reads. Second, unaligned reads with 3’-CCA were randomly assessed by blasting to 3D7 genome to ensure most reads with 3’-CCA were mapped in prior operations. After alignment, reads with the same UMI and mapping coordinates were de-duplicated using UMI_tools (v1.1.2)^71^.

tRNA composition was determined by the ratio of read counts per tRNA to the read counts of all tRNAs found by htseq-count with a minimum alignment quality of 10 (v0.9.1)^75^.

The determination for tRNA aminoacylation level is as follows: an aligned read is considered CCA-ending(charged) if it aligns with no mismatches to the tRNA’s 3’ end. Consequently, if the read aligns to the 3’end but only ends in -CC, it is considered uncharged. Aminoacylation level were determined as the percentage of CCA-ending reads over the sum of the CCA-ending and CC-ending reads, adjusted to the recovered ratio of the two spike-in RNA oligos.

Fractions of mutation signature were calculated as 1-(coverage of reference sequence at position n/ total coverage at position n+1).

#### mRNA library read processing

mRNA libraries were sequenced on an Illumina Nextseq 550 platform with a single-end read length of 120 bp or 150bp (for the polysome profiles). Base calling was performed with Generate FASTQ (v2.0.1). Sequence reads were first filtered with Trimmomatic (v0.38) using the default setting^76^. The trimmed reads were then aligned to 3D7 genome (v. PlasmoDB-46) with HISAT2 (v2.2.1) using default setting and read counts were obtained by htseq-count (v0.9.1)^75,77^.

### Amino acid starvation assay

Parasite cultures were synchronized with two consecutive cycles of sorbitol and percoll gradient treatment before the start of the assay. All starvation assays used parasites at 32-36 hpi (pre-segmented) maintained at 3% hematocrit and >5% parasitemia. The parasites were washed twice with and resuspended in either AA-free RPMI (US biological) or complete RPMI (US biological) supplemented with 0.5% AlbumaxII and 20µg/ml hypoxanthine, pH 7.2. Parasites were incubated for 6 h and harvested immediately afterwards for the preparation of tRNA library and quantitative proteomics analysis. 70 nM of halofuginone (Calbiochem) or 382 nM of L-isoleucine (Sigma) were also added where applicable.

### Quantitative proteomics analysis

#### Sample preparation

Cell pellets were solubilized with 100 µL of 8M urea, 0.2% ProteaseMAX (Promega) and 5% acetonitrile (ACN) in 100 mM Tris-HCl, pH 8.0 with 1 µL of 100x protease inhibitor cocktail (Roche) during sonication in water bath for 10 min followed by probe sonicated with VibraCell probe (Sonics & Materials, Inc.) for 20 s, with pulse 2/2, at 20% amplitude. Following centrifugation at 10,000 g for 10 min at 4°C, the supernatant was transferred to a new tube and the remaining pellet was supplemented with 50 µL of 1M urea in 100 mM Tris-HCl with 0.2% ProteaseMAX and sonicated in water bath for 10 min. Protein concentration in the combine supernatant was measured by BCA assay (Thermofisher Scientific). An aliquot of 25 µg samples was transferred to a new tube and equalized with 50 mM Tris-HCl to a total volume of 100 µL. Proteins were reduced with addition of 1 µL of 500 mM dithiothreitol (Sigma) at 25°C for 1 h and alkylated with 3 µL of 500 mM iodoacetamide for 1 h at room temperature in the dark and quenched with 2 µL of dithiothreitol for 10 min. Proteolytic digestion was started with 2 µL of 0.5 µg/µL Lys-C (Wako, Japan) at 37°C for 3 h with shake at 450 rmp and supplemented with 300 µL of 100 mM Tris-HCl, pH 8.0 before adding 4 µL of 0.5 µg/µL sequencing grade trypsin (Promega) and incubated over night at 37°C. The digestion was stopped with 20 µL cc. formic acid, incubating the solutions at RT for 5 min. The sample was cleaned on a C18 Hypersep plate with 40 µL bed volume (Thermofisher Scientific), dried using a vacuum concentrator (Eppendorf). Biological samples were labeled with TMT-10plex reagents in random order adding 200 µg TMT-reagent in 30 µL dry ACN to each digested sample resolubilized in 70 µL of 50 mM triethylammonium bicarbonate (TEAB) and incubating at room temperature (RT) for 2 h. The labeling reaction was stopped by adding 11 µL of 5% hydroxylamine and incubated at RT for 15 min before combining them in one vial.

#### Liquid Chromatography-Tandem Mass Spectrometry Data Acquisition

Peptides were reconstituted in solvent A and approximately, 2 µg samples injected on a 50 cm long EASY-Spray C18 column (Thermofisher Scientific) connected to an Ultimate 3000 nanoUPLC system (Thermofisher Scientific) using a 120 min long gradient: 4-26% of solvent B (98% acetonitrile, 0.1% FA) in 120 min, 26-95% in 5 min, and 95% of solvent B for 5 min at a flow rate of 300 nL/min. Mass spectra were acquired on a Q Exactive HF hybrid quadrupole orbitrap mass spectrometer (Thermofisher Scientific) ranging from m/z 375 to 1700 at a resolution of R=120,000 (at m/z 200) targeting 5×106 ions for maximum injection time of 80 ms, followed by data-dependent higher-energy collisional dissociation (HCD) fragmentations of precursor ions with a charge state 2+ to 8+, using 45 s dynamic exclusion. The tandem mass spectra of the top 18 precursor ions were acquired with a resolution of R=60,000, targeting 2×105 ions for maximum injection time of 54 ms, setting quadrupole isolation width to 1.4 Th and normalized collision energy to 33%.

#### Data analysis

Acquired raw data files were analyzed using Proteome Discoverer v2.5 (Thermofisher Scientific) with Sequest HT search engine against Plasmodium falciparum PF3D7 protein database (SwissProt). A maximum of two missed cleavage sites were allowed for full tryptic digestion, while setting the precursor and the fragment ion mass tolerance to 10 ppm and 0.02, respectively. Carbamidomethylation of Cys was specified as a fixed modification, while TMT6plex on Lys and N-termini, oxidation on Met as well as deamidation of Asn and Gln were set as dynamic modifications. Initial search results were filtered with 5% FDR using Percolator node in Proteome Discoverer. Quantification was based on the TMT-reporter ion intensities. Differential quantitation analysis was performed by DESeq2^78^.

### Multiplication rate assay

Parasite cultures were synchronized with two consecutive cycles of sorbitol and percoll gradient treatment before the start of the assay. Parasites at 8-12 hpi or 32-36 hpi were collected for the assay. The parasites were washed twice in AA-free RPMI and then resuspended at 2% hematocrit and 1% parasitemia in different conditions including in complete medium, AA-free medium and AA-free medium supplied with varying concentration of isoleucine (382 μM, 40 μM, 20 μM, 10 μM, 5 μM, 2.5 μM) in a 96-well plates. After 6h of incubation, the starved cultures were re-supplemented with complete medium until all parasites had entered a new invasion cycle. Microscopy assessment was used to ensure all parasites were either in ring or trophozoite stage. Samples were then harvested for flow cytometry analysis.

### Flow cytometry analysis

Samples were harvested for flow cytometry analysis at 0h, 6h, 24h and 48h after plate setup for 6-10 hpi parasites and at 0h, 6h, 12h, 18h and 32h for 32-36hpi parasites. 20µL of sample per timepoint were harvested and stained for 30 min at 37°C with Hoechst (Invitrogen) and DHE (Sigma) at a final concentration of 20 µg/ml and 10 µg/ml respectively. Stained cells were diluted in PBS to a final hematocrit of 0.2% prior to analysis. At least 3000 Hoechst+/DHE+ events/sample were acquired using a FACSVerse (BD Biosciences). Data was analyzed as described previously using FlowJo (Version 10.6.2)^79^.

### Polysome fractionation

Polysome purification was adopted from a protocol previously reported but with major modifications^80^.

Cultures were expanded to 150 mL at 3% hematocrit and ∼10% parasitemia and were synchronized with two consecutive cycles of sorbitol and percoll gradient treatment before the start of the assay. When the culture reached 32-36 hpi, infected RBCs were enriched with magnetic separation using XS column (Miltenyi Biotec). After the enrichment, the cell pellet was equally split and cultured either in AA-free RPMI or complete RPMI for 6h.

After the incubation, cycloheximide was added to a final concentration of 100 µg/mL to the cultures. The cultures were briefly shaken and immediately incubated on ice for 10 min. Afterwards, the cells were centrifuged and washed once with 10 ml ice-cold PBS supplemented with 100 µg/ml cycloheximide. Equal volume of ice-cold 2x lysis buffer (800 mM potassium acetate, 50 mM potassium HEPES, pH 7.2, 30 mM magnesium acetate, 10 µg/mL cycloheximide, 2 mM DTT, 2 mM AEBSF, 40 U/mL rRNasin (Promega), 1% sodium deoxycholate, 2% Igepal CA-630) was added to the pellet and the pellet was gently resuspended. The resuspended pellet was incubated on ice for 5 min with a brief vortex every minute. The sample was then centrifuged at 17000 g for 2 min to clear the lysate. 50 µL of the lysate was separated for total RNA extraction.

A 10% to 50% sucrose gradient solution in gradient buffer (20 mM HEPES pH 7.6, 100 mM KCl, 5 mM MgCl_2_) was prepared on a Gradient Station platform (Biocomp Instruments). 450 µl of the lysate was carefully layered on top of the sucrose gradient and centrifuged at 35000 rpm (210000g) for 2h on a pre-chillled SW40 Ti rotor with the OptimaXL-100K ultracentrifuge system (Beckman Coulter). After the centrifugation, the gradient was fractionated into 25 fractions on a Gradient Station platform (Biocomp Instruments). RNA and proteins were extracted from the fractions using Trizol reagent (Sigma) as per manufacturer’s instruction and the RNA from the polysomal fractions (≥3 associated ribosomes, fractions 15-25) were pooled.

### Immunoblotting

Western blot was performed with proteins purified from the polysome fractionation. Purified proteins were solubilized in Laemmli buffer (Bio-Rad) with 2% β-mercaptoethanol and heated at 95°C for 10 min. The proteins were resolved in a criterion TGX Stain-Free gel (Bio-Rad) with Tris/Glycine/SDS running buffer (Bio-Rad). The resolved proteins were transferred to nitrocellulose membrane in Tris/glycine buffer (Bio-Rad). The membrane was blocked with 5% skim milk, 2% BSA in PBS at 4°C overnight. The membrane was incubated with 1:2000 anti-RPS14 antibody (abcam, #ab174661) in 2% skim milk, 2% BSA in PBS for 2h at room temperature. The membrane was washed thrice with 1x TBS, 0.1% Tween 20 and then incubated for 30 min with 1:5000 HRP-conjugated anti-rabbit IgG (Amersham, #NA934). Chemiluminescent signal was detected with a ChemiDoc imager (Bio-Rad) after developing with WesternBright ECL kit (advansta).

### Orthologue analysis

All orthologues to the isoleucine-rich genes of *P. falciparum* were obtained from OMA orthology database using the OMA group fingerprint (Dec 2021 release) ^24^. A cut-off of having at least 100 orthologues was applied to consider a *P. falciparum* protein as functionally conserved and used for analysis.

### Kozak Similarity score (KSS)

Calculation of KSS was inspired by a previous report^24^. Bit scores for each nucleotide from position -3 to +4 of the annotation start codon was generated from all genes encoding ribosomal protein and used as a reference. The KSS of a gene was calculated as the sum of all the bit scores from these positions relative to the maximum possible score summed from these positions.

### Statistical analysis

Two tailed student’s t-test, two tailed Mann-Whitney U test and Fisher’s exact test were used for statistical analyses wherever specified. A p-value of <0.05 was used as the cut off for statistical significance. Multiple testing were corrected with Benjamini-Hochberg procedure, with a false discovery rate <0.05 used as the cut off for statistical significance.

### Gene ontology (GO) enrichment analysis

GO term analysis was performed using the PlasmoDB GO term pipeline using “biological process” as the ontology and with a p-value cutoff of ≤ 0.05. FDR<0.05 was used as cut-off for significantly enriched pathways.

## Acknowledgement

We thank all members of the Larsson laboratory and the Ribacke laboratory for their constructive comments and feedback. We thank Pan Tao and Addgene for providing us the demethylase plasmids and E. Strandback, H.A. Korsah and T. Nyman in the Protein Science facility of Karolinska Institutet for assistance in synthesizing the recombinant proteins. Protein identification and quantification were carried out by the Proteomics Biomedicum core facility, Karolinska Institutet (https://ki.se/en/mbb/proteomics-biomedicum). The authors acknowledge the support of the Freiburg Galaxy Team: Person *X* and Björn Grüning, Bioinformatics, University of Freiburg (Germany) funded by the Collaborative Research Centre 992 Medical Epigenetics (DFG grant SFB 992/1 2012) and the German Federal Ministry of Education and Research BMBF grant 031 A538A de.NBI-RBC. This work was supported by a grant from Knut and Alice Wallenberg Foundation (2017.0055.) to M.W. and U.R, a grant from the Swedish Research Council (2021-03141) to S.CL.C. and M.W, a grant from Tore Nilsons Stiftelse För Medicinsk Forskning (2017-00532) to S.CL.C., S.CL.C. was also supported by a postdoctoral fellowship grant from Svenska Sällskapet för Medicinsk Forskning, Q.L. was supported by a scholarship from Chinese Scholarship Council and a doctoral (KID #2018-01037) grant from Karolinska Institutet.

## Author contributions

S.CL.C. conceived the study. S.CL.C. and Q.L designed the experimental setups. S.CL.C. and Q.L. optimized the tRNAseq protocol. Q.L collected the samples and generated the sequencing libraries. S.CL.C. and Q.L. performed all the data analyses unless otherwise specified. L.V. performed the flow cytometry experiment and its data analyses. A.V. performed the quantitative proteomic experiment and its data analyses. S.CL.C. performed the polysome profiling and its data analyses. C.S., U.R. and O.L. provided critical inputs for the data analyses. S.CL.C. and Q.L. wrote an initial draft of the manuscript and generated all the figures with contributions from all authors. All authors edited the manuscript. The overall project was supervised by S.CL.C., O.L. and M.W.

## Ethics declaration

The authors declare no competing interests.

