## Extended data file for "Amino acid usage and tRNA expression underlies proteome adaptation in *Plasmodium falciparum*"

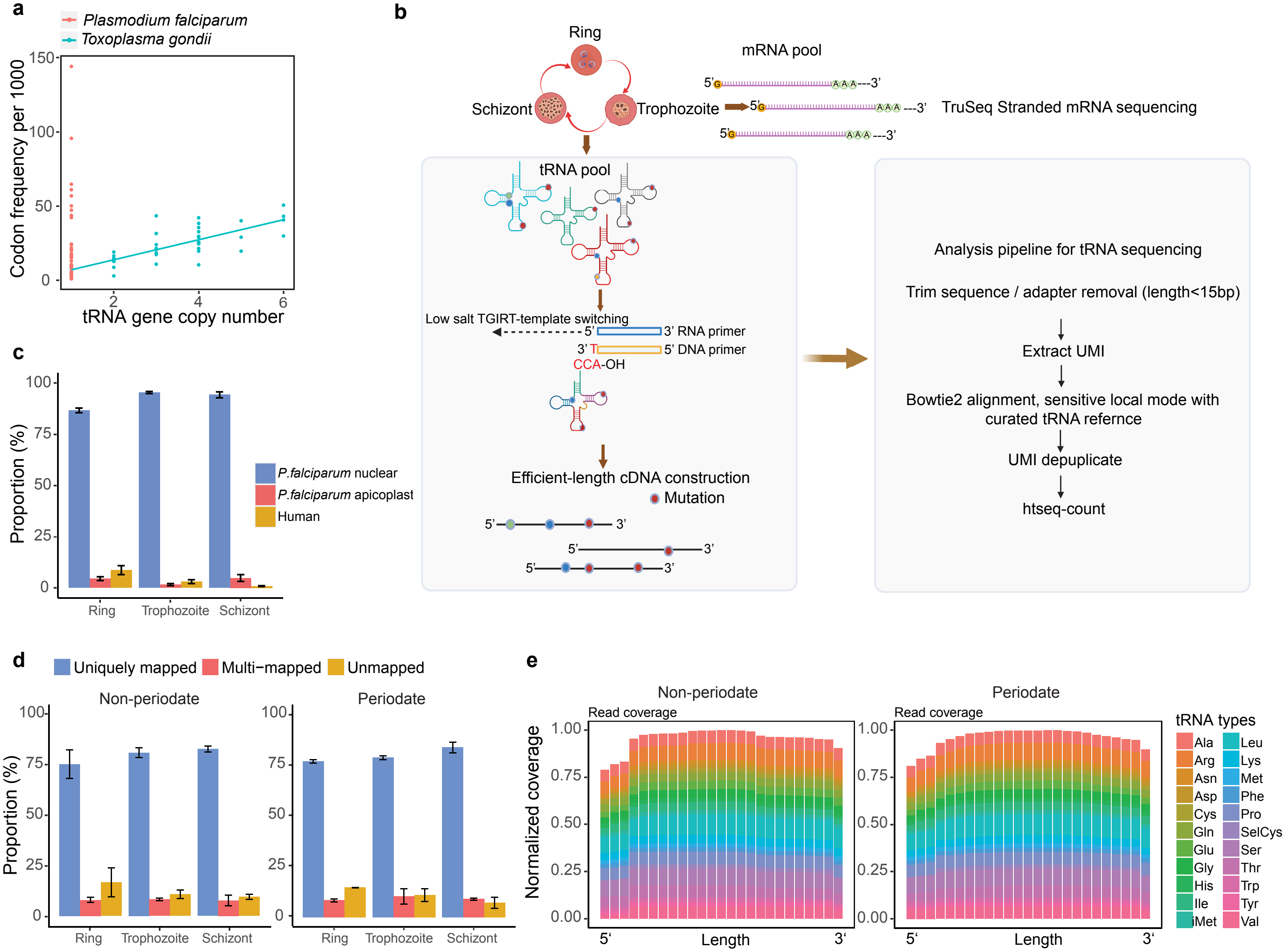
a
Plasmodium falciparum
Toxoplasma gondii
Codon frequency per 1000
tRNA gene copy number
b
Ring
Schizont
Trophozoite
mRNA pool
TruSeq Stranded mRNA sequencing
tRNA pool
Low salt TGIRT-template switching
3' RNA primer
5' DNA primer
CCA-OH
Efficient-length cDNA construction
Mutation
Analysis pipeline for tRNA sequencing
Trim sequence / adapter removal (length<15bp)
Extract UMI
Bowtie2 alignment, sensitive local mode with curated tRNA refernce
UMI depuplicate
htseq-count
c
Proportion (%)
P.falciparum nuclear
P.falciparum apicoplast
Human
Ring
Trophozoite
Schizont
d
Uniquely mapped
Multi−mapped
Unmapped
Non-periodate
Periodate
Proportion (%)
Ring
Trophozoite
Schizont
e
Non-periodate
Periodate
Read coverage
Normalized coverage
Length
tRNA types
Ala
Arg
Asn
Asp
Cys
Gln
Glu
Gly
His
Ile
iMet
Leu
Lys
Met
Phe
Pro
SelCys
Ser
Thr
Trp
Tyr
Val

Extended data Fig.1

**tRNAome analysis of *P. falciparum*.**

**a,** Relationship between tRNA gene copy numbers and codon usage in *T. gondii* and *P. falciparum.* **b,** Left panel: Schematic showing tRNA sequence library generation using TGIRT. A DNA/RNA duplex with a T overhang complementary to the terminal adenosine of mature tRNAs was used for the template-switching reaction. The 3' adapter was ligated after the cDNA was synthesized. Right panel: Strategy for read processing of the tRNA sequencing libraries. **c,** The origin of all reads mapped to tRNAs throughout the IDC. Error bars represent the standard deviation of N=3 biological replicates. **d,** Bowtie2 alignment-statistics for tRNA-sequencing data generated from non-periodate (untreated, N=3) and periodate-treated (N=3) libraries. Error bars represent standard deviation from N=3 biological replicates. **e,** Representative scaled 5'-to-3' sequence coverage plots spanning all nuclear tRNAs. The full-length tRNA was divided into 25 bins with each bin representing 4% of the tRNA length, as shown on the x-axis. The Y-axis value for each tRNA was normalized to the bin with the maximum coverage for each respective tRNA. Left panel: samples without periodate treatment. Right panel: sample treated with periodate.

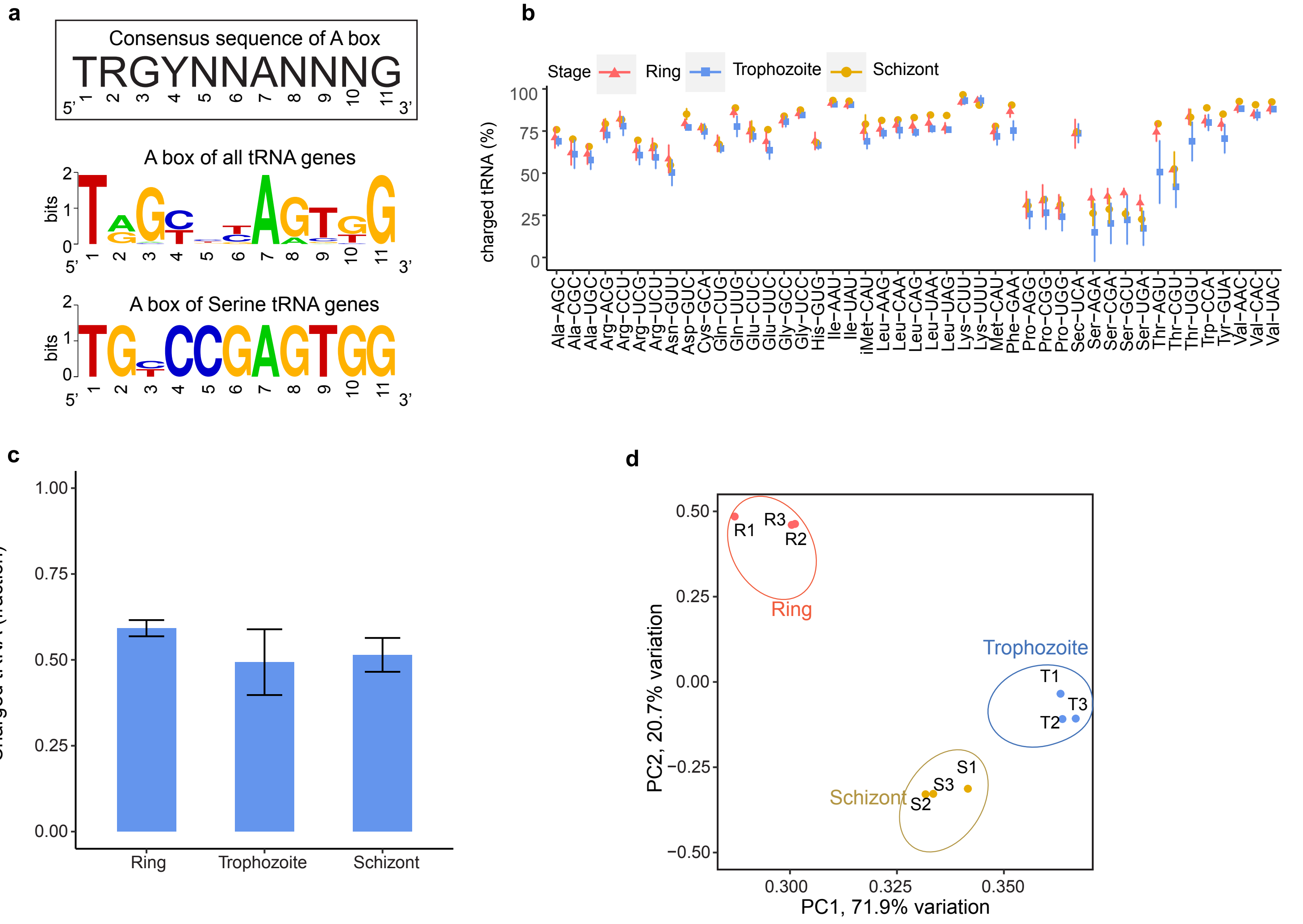


Extended data Fig.2

**Characteristics of the anticodon and codon pools.**

**a,** Motif logo of the A-box in all, except Ser-tRNAs, *P. falciparum* tRNAs (upper), and of all Ser-tRNAs (lower). With a G-to-Y substitution at the third position, the A-box of Ser-tRNAs diverge from the eukaryotic consensus (boxed). **b,c,** tRNA charging throughout the IDC. Relative charge of (**b**) individual tRNA and **(c)** total tRNA throughout the IDC. Charged tRNAs are represented by reads with 3' -CCA ends after periodate oxidation and beta-elimination reactions. Error bars represent the standard deviation of N=3 biological replicates for Ring and Trophozoite, Schizont: N=2. All apicoplast tRNAs are excluded **d,** Principal component analysis of the coding transcriptomes throughout the IDC.

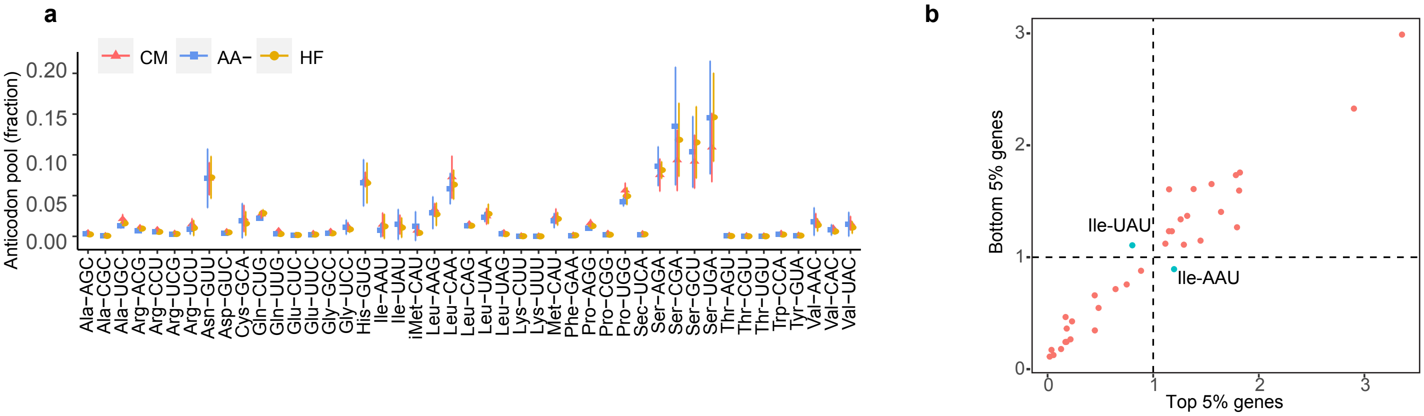


Extended data Fig.3

**Limited changes in the tRNAome after acute amino acid depletion.**

**a,** Changes in tRNA composition after 6 hours of treatment in late stage parasite 32-36 h.p.i. (CM: complete medium; AA-: AA starvation; and HF: halofuginone (70 nM) and AA starvation). Error bars represent the SD of N=3 biological replicates. **b,** The relative isoacceptor usage of the 5% most highly expressed coding genes (x-axis, n=258) and the 5% least expressed coding genes (y-axis, n=258). Relative isoacceptor usage for each tRNA was calculated as the fraction of all codons encoding a specific amino acid that is decoded by an isoacceptor relative to the fraction of all isoacceptors for a specific amino acid that are equally used. The most highly expressed genes preferred Ile-AAU, while lowly expressed genes preferred Ile-UAU.

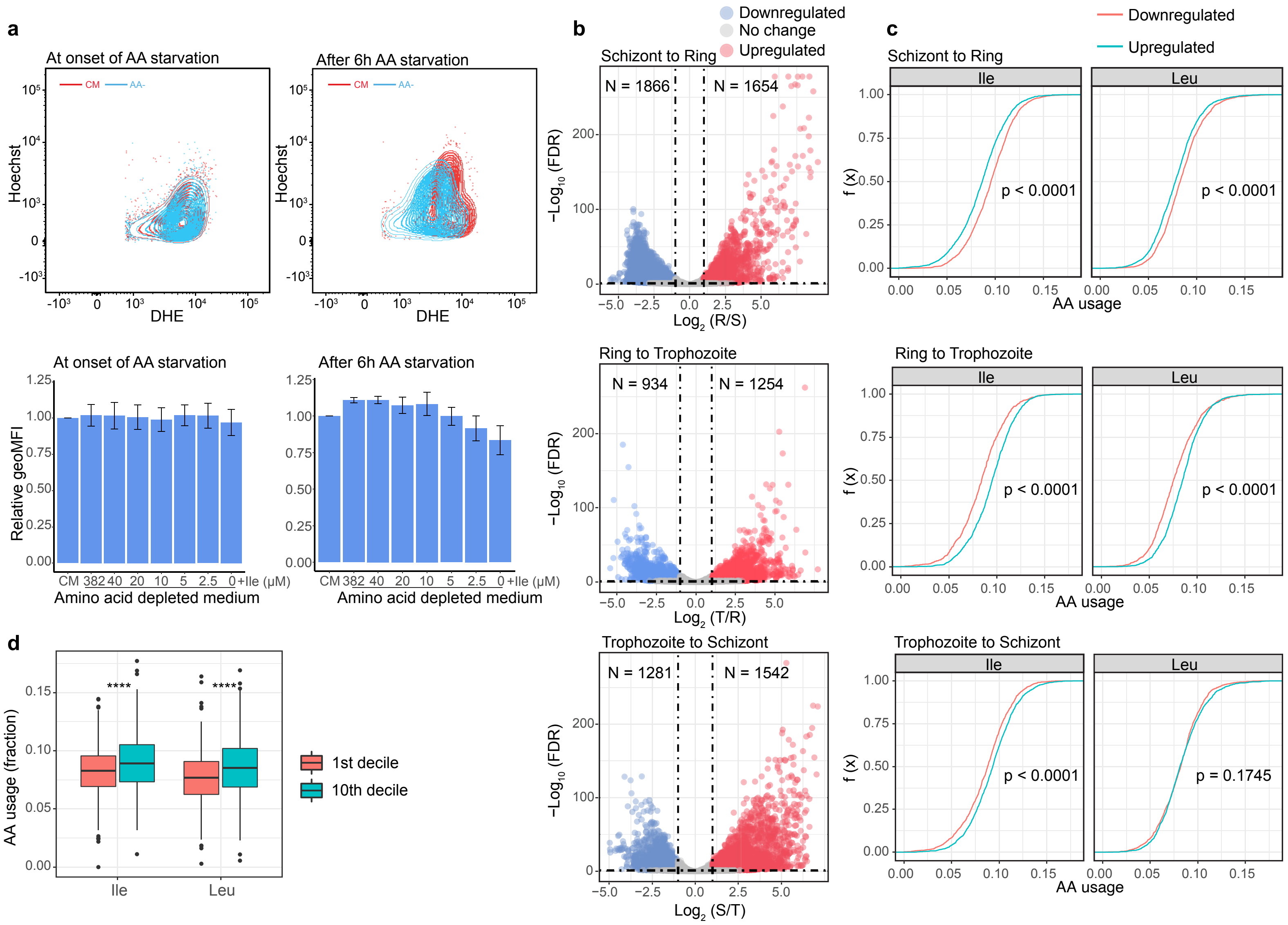

a
At onset of AA starvation
After 6h AA starvation
CM
AA-
Hoechst
DHE
Relative geoMFI
CM 382 40 20 10 5 2.5 0 +Ile (μM)
Amino acid depleted medium
b
Downregulated
No change
Upregulated
Schizont to Ring
N = 1866
N = 1654
−Log10 (FDR)
Log2 (R/S)
Ring to Trophozoite
N = 934
N = 1254
Log2 (T/R)
Trophozoite to Schizont
N = 1281
N = 1542
Log2 (S/T)
c
Downregulated
Upregulated
Schizont to Ring
Ile
Leu
f (x)
p < 0.0001
AA usage
Ring to Trophozoite
p < 0.0001
p < 0.0001
Trophozoite to Schizont
p < 0.0001
p = 0.1745
d
AA usage (fraction)
****
****
1st decile
10th decile
Ile
Leu

Extended data Fig.4

**Stage progression is isoleucine sensitive and is delayed by acute amino acid deprivation**

**a,** FACS plots showing the fluorescence intensity of dihydroethidium (DHE) and Hoechst DNA staining in parasites (upper left) at the beginning and (upper right) after 6 hours of AA starvation (AA-) compared to parasites grown in complete medium (CM). Both DHE and Hoechst stainings were reduced in AA- parasites after 6 hours. The geometric mean of the Hoechst staining intensity in parasites (lower left) at the beginning and (lower right) after 6 hours of AA starvation relative to that in CM parasites is shown. Supplementing isoleucine to AA- parasites rescued the reduction in DNA content in a dose-dependent manner. Error bars represent the SD for N=3 biological replicates. **b,** DESeq2 differential gene expression analyses of the transcriptomes during stage progression from (upper) the schizont-to-ring stage, (middle) the ring-to-trophozoite stage, and (below) the trophozoite-to-schizont stage. The horizontal and vertical dotted lines show the applied adjusted p-value (<0.05) and fold change (2-fold) thresholds, respectively. N=3 biological replicates. **c,** Empirical cumulative distribution function (CDF) plot of isoleucine and leucine demand in up- (cyan) and down-regulated (red) genes during stage progression. The p-values of Mann–Whitney U tests are indicated. **d,** Isoleucine and leucine usage in the 10% of genes with the most variable expression (the 10th decile) and the 10% of genes with the least variable expression (1st decile) throughout the IDC. Variability was defined by the coefficient of variation in expression (CV of TPM) throughout the IDC.

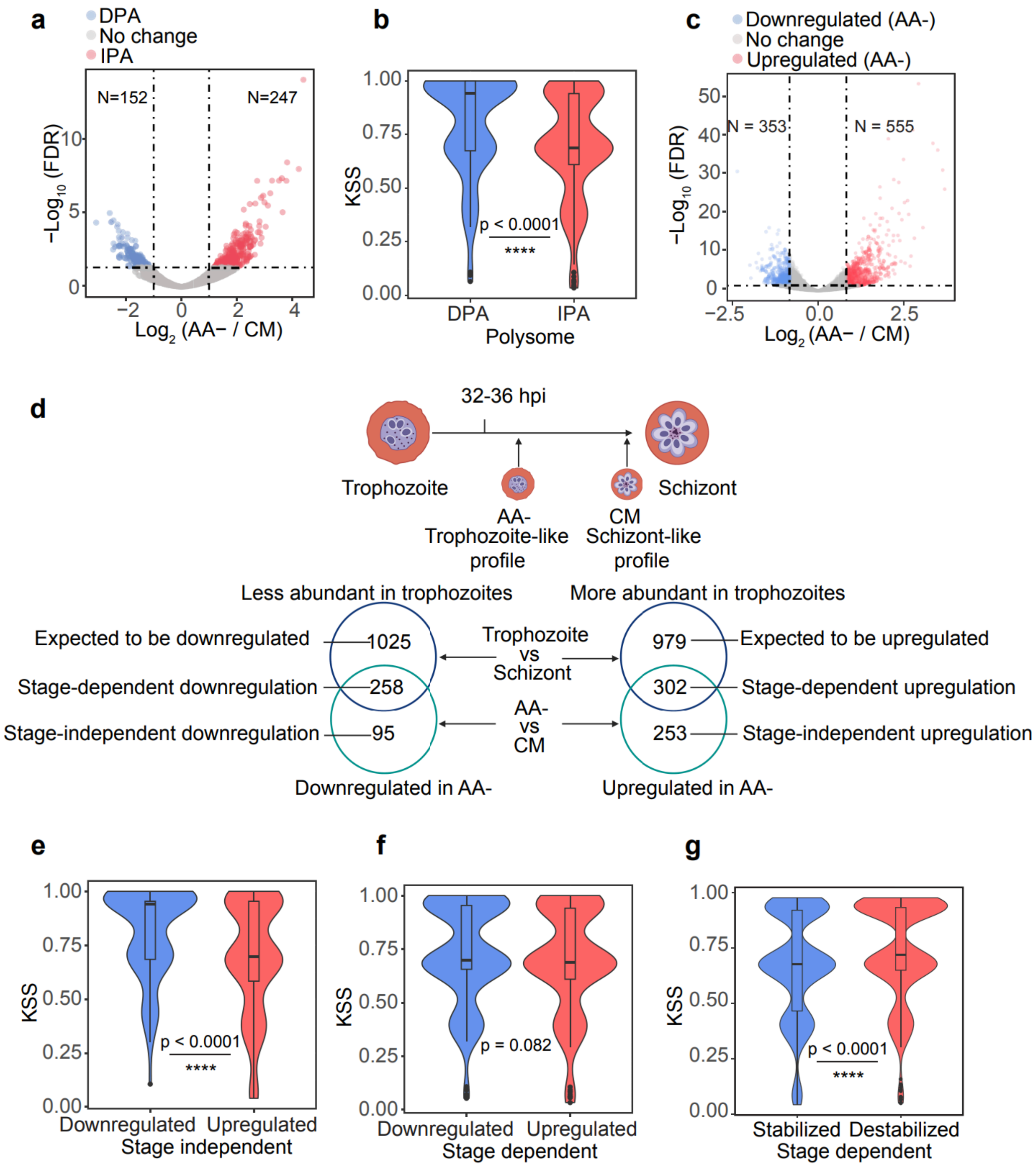
a
DPA
No change
IPA
N=152
N=247
−Log10 (FDR)
Log2 (AA− / CM)
b
KSS
p < 0.0001
****
DPA
IPA
Polysome
c
Downregulated (AA-)
No change
Upregulated (AA-)
N = 353
N = 555
−Log10 (FDR)
Log2 (AA− / CM)
d
32-36 hpi
Trophozoite
Schizont
AA-
Trophozoite-like
profile
CM
Schizont-like
profile
Less abundant in trophozoites
More abundant in trophozoites
Expected to be downregulated
1025
Trophozoite
vs
Schizont
979
Expected to be upregulated
Stage-dependent downregulation
258
302
Stage-dependent upregulation
Stage-independent downregulation
95
AA-
vs
CM
253
Stage-independent upregulation
Downregulated in AA-
Upregulated in AA-
e
KSS
p < 0.0001
****
Downregulated
Upregulated
Stage independent
f
KSS
p = 0.082
Downregulated
Upregulated
Stage dependent
g
KSS
p < 0.0001
****
Stabilized
Destabilized
Stage dependent

Extended data Fig.5

**Increased ribosome loading stabilizes transcripts and reprograms the transcriptome**

**a,** DESeq2 differential gene expression analysis for polysome-associated mRNA after AA starvation. The horizontal and vertical dotted lines show applied adjusted p-value (<0.05) and fold changes (2-fold) cutoffs, respectively. **b,** Kozak similarity scores (KSS, using ribosomal genes as reference) in transcripts with increased polysome association (IPA) or decreased polysome association (DPA) after amino acid depletion. Transcripts with increased polysome association had lower scores, indicating an initiation context less similar to that of the highly translated ribosome encoding mRNAs. The p-value from a Mann– Whitney U test is indicated. **c,** DESeq2 differential gene expression analysis of the transcriptome after AA starvation. The horizontal and vertical dotted lines show applied adjusted p-value (<0.05) and fold change (2-fold) cutoffs, respectively. N=3 biological replicates. **d,** An overview of the strategy used to categorize stage-dependent and stage-independent changes in the transcriptome. The transcriptome of the AA-starved parasites (AA-) was expected to most closely resemble a trophozoite-like transcriptional profile due to delayed stage progression (upper). Transcripts that were more abundant in trophozoites were expected to be upregulated in AA- parasites and vice versa, as illustrated in the Venn diagram. **e,f,** The KSS of stage-independent (**e**) and stage-dependent (**f**) up- and downregulated genes. Only stage-independent changes show a disparity in KSS between the up- and downregulated genes similar to that in the polysome fractions, supporting a positive feedback loop. The p-values from Mann–Whitney U tests are indicated. **g,** The KSS of transcripts that remained stable after AA starvation but were expected to be downregulated (and therefore stabilized) or upregulated (and therefore destabilized) according to the categorization shown in (**d**). Statistical significance was assessed using Mann-Whitney U tests in (e-g). Resulting p-values are indicated.

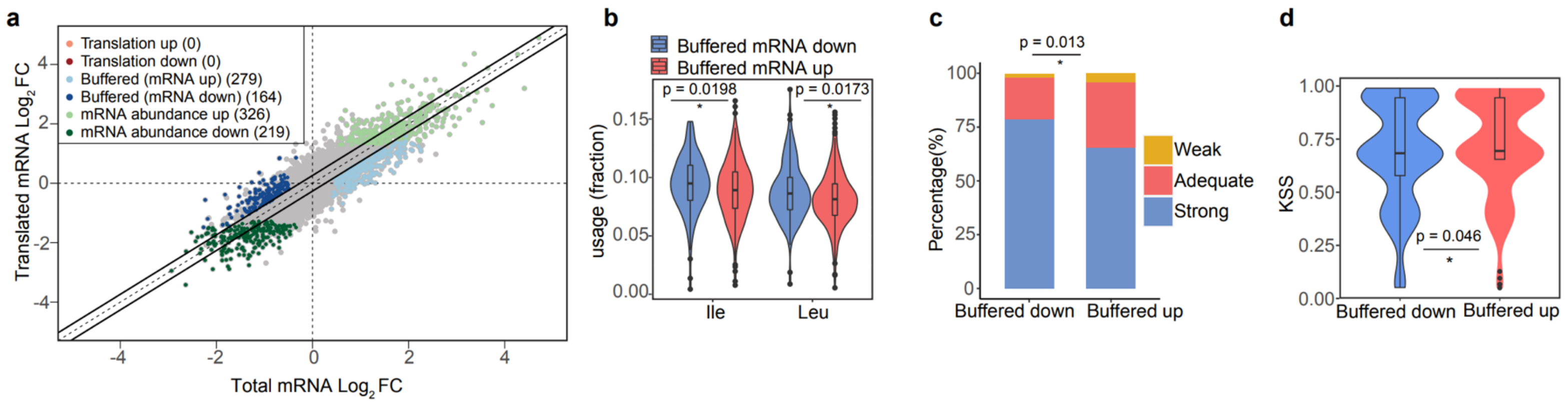


Extended data Fig.6

**Anota2seq analysis reveals an abundance of translationally "buffered" transcripts**

**A,** Anota2seq analysis of the polysome-profiling dataset using rlog normalization with default parameters. Anota2seq performs analysis of multiple gene expression modes. Genes classified as buffered (mRNA down) correspond to those with a relative increase in ribosome loading but with no transcript-stabilizing effect. N=3 biological replicates. **b-d**, The differences in (**b**) isoleucine and leucine usage, the Kozak context (**c**), and KSS (**d**) in "buffered (mRNA up)" and "buffered (mRNA down)" transcripts.

Kozak context: strong, RnnATGR; adequate, RnnATGY/YnnATGR; and weak, YnnATGY (ATG= start codon).

Statistical significance was assessed using Mann-Whitney U tests in (b,d); and Fisher's exact test for the Kozak context analysis in (c). Resulting p-values are indicated.

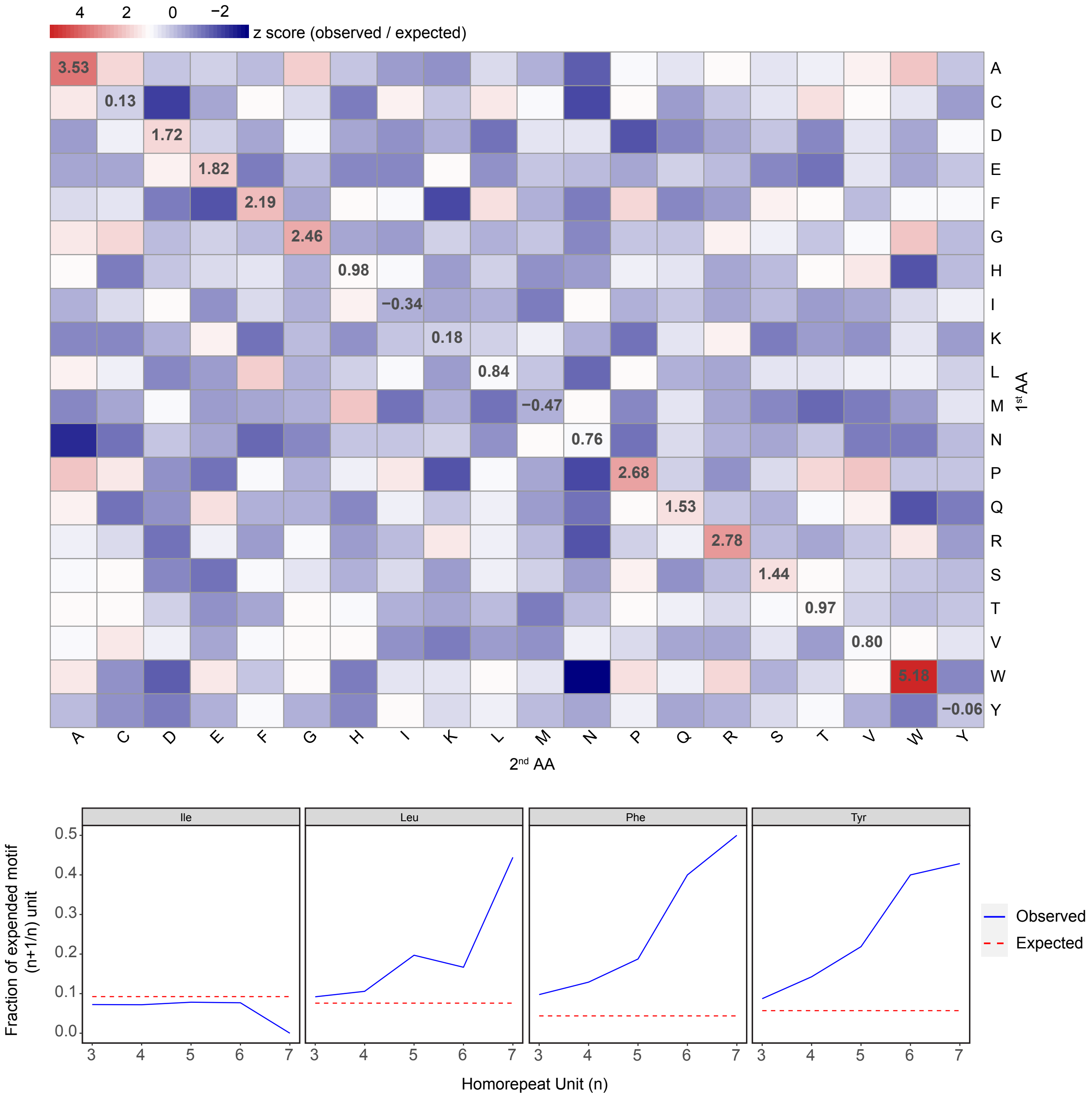


Extended data Fig.7

**The expansion of Ile homorepeats is under purifying selection.**

(Upper) Heatmap showing the frequency of di-amino acid motifs relative to their expected frequency. Values of (observed/expected frequency) were calculated for each di-amino acid motif, and all values were Z-transformed to generate the heatmap. (Lower) The fraction of homorepeat motifs retained with successive expansion in residue number (n) in four types of hydrophobic amino acid. The dotted lines represent the expected frequency based on the amino acid usage. Four of the most frequently used hydrophobic amino acids were analysed.
